# Synovial macrophage diversity and activation of M-CSF signaling in post-traumatic osteoarthritis

**DOI:** 10.1101/2023.10.03.559514

**Authors:** Alexander J. Knights, Easton C. Farrell, Olivia M. Ellis, Michelle J. Song, C. Thomas Appleton, Tristan Maerz

**Author notes:** Corresponding author; Rm 2017, Biomedical Science Research Building, 109 Zina Pitcher Pl, Ann Arbor, 48109.

## Abstract

**Objective:** Synovium is home to immune and stromal cell types that orchestrate inflammation following a joint injury; in particular, macrophages are central protagonists in this process. We sought to define the cellular and temporal dynamics of the synovial immune niche in a mouse model of post-traumatic osteoarthritis (PTOA), and to identify stromal-immune crosstalk mechanisms that coordinate macrophage function and phenotype.

**Design:** We induced PTOA in mice using a non-invasive tibial compression model of anterior cruciate ligament rupture (ACLR). Single cell RNA-seq and flow cytometry were used to assess immune cell populations in healthy (Sham) and injured (7d and 28d post-ACLR) synovium. Characterization of synovial macrophage polarization states was performed, alongside computational modeling of macrophage differentiation, as well as implicated transcriptional regulators and stromal-immune communication axes.

**Results:** Immune cell types are broadly represented in healthy synovium, but experience drastic expansion and speciation in PTOA, most notably in the macrophage portion. We identified several polarization states of macrophages in synovium following joint injury, underpinned by distinct transcriptomic signatures, and regulated in part by stromal-derived macrophage colony-stimulating factor signaling. The transcription factors Pu.1, Cebpα, Cebpβ, and Jun were predicted to control differentiation of systemically derived monocytes into pro-inflammatory synovial macrophages.

**Conclusions:** We defined different synovial macrophage subpopulations present in healthy and injured mouse synovium. Nuanced characterization of the distinct functions, origins, and disease kinetics of macrophage subtypes in PTOA will be critical for targeting these highly versatile cells for therapeutic purposes.

## INTRODUCTION

Arthritis encompasses a host of disease manifestations in joints, characterized in part by aberrant crosstalk between stromal and immune cells, with a central unifying feature of disease being inflammation. Inflammation is driven by a diverse and highly plastic set of resident and infiltrating immune cells that can speciate into distinct functional phenotypes and engage in reciprocal crosstalk with stromal cells. While a physiological inflammatory response is integral to restoring tissue homeostasis after insults such as trauma or infection, aberrant immune cell responses underpin chronic inflammation and its associated pathological effects in many tissue contexts. For instance, early studies of diet-induced obesity in mice showed recruitment of pro-inflammatory macrophages to adipose tissue from circulating monocytes^1, 2^. In systemic sclerosis, interactions between distinct macrophage subsets and fibroblasts drive them to adopt pro-inflammatory and pro-fibrotic phenotypes, respectively, that underpin disease progression^3^. Specifically, osteopontin produced by pathologic *SPP1+* macrophages promotes production of type I collagen by skin fibroblasts^4^. Fibroblasts in sclerotic lesions produce high levels of TLR4 that drive persistent fibrosis^5^.

The synovium is a heterogenous connective tissue in joints, populated with immune cells, vascular and lymphatic endothelial cells, fibroblasts, adipocytes, and other mesenchymal-lineage stromal cells^6^. Synovium is increasingly recognized as a principal regulator of intra-articular inflammation in osteoarthritis (OA), given its highly vascularized nature and abundance of resident immune cells compared to tissues such as meniscus and cartilage. Synovial inflammation, now appreciated as a pathologic driver of OA^7-9^, is characterized by expansion of resident synovial fibroblasts and immune cells as well as influx of systemically derived immune cells; however, we have a limited understanding of the cellular, molecular, and temporal mechanisms of these interactions, and how they differ or correspond across forms of arthritis.

The specific roles of distinct activated immune cell populations in orchestrating synovial inflammation have been well characterized in the context of rheumatoid arthritis (RA) ^10, 11^, providing valuable insights of disease pathogenesis and advancing therapeutic efficacies^12^. Our understanding of pathologic immune cell landscape in post-traumatic OA (PTOA) synovium and its contribution to inflammation is only in its infancy, however.

Whether macrophage expansion is a necessary mechanism to restore homeostasis or a deleterious pathological manifestation is still controversial. Macrophage ablation in mice by clodronate liposomes yielded generally favorable OA outcomes, including reduced osteophyte formation^13, 14^, however pharmacogenetic macrophage depletion using the MaFIA mouse resulted in systemic inflammation, exacerbated immune influx into the joint, with no attenuation of OA severity^15^. Indeed, synovial macrophages play critical physiological functions such as the timely clearance of apoptotic cells, which is impaired in OA^16^. In a model of inflammatory arthritis, Huang et al showed that resolution requires the suppression of pro-inflammatory, systemically-derived F4/80^hi^ MHC Class II^+^ macrophages and that macrophages with a resident-like phenotype promote inflammatory resolution^17^. As such, dissecting macrophage identity, origin, and function in arthritis remains critical to better understanding disease etiology.

In this study, we employed a non-invasive anterior cruciate ligament rupture (ACLR) model of PTOA that recapitulates traumatic joint injury^18, 19^ to dissect the dynamic immune cell environment in PTOA synovium, with a focus on the early post-injury period. Importantly, the ACLR model avoids the need for surgery and its confounding effects on synovial inflammation. We utilized single cell RNA sequencing (scRNA-seq) and flow cytometry to identify distinct immune cell types in healthy and PTOA synovia, with a focus on macrophages, and identified potential stromal-derived crosstalk mechanisms responsible for emergence of pro-inflammatory macrophage subtypes.

## METHODS

### Mice

Male and female mice on a C57Bl/6 background were used for all experiments. Mice were housed in ventilated cages of up to five animals, given chow food and water, on a 12 h light/dark cycle. At time of injury, mice were 12-14 weeks old, and were euthanized by CO2 asphyxia. We induced PTOA using a previously reported and characterized model involving non-invasive rupture of the anterior cruciate ligament (ACLR) ^20^. Sham mice were given anesthesia and analgesic, but not subjected to injury. All protocols were conducted in accordance with approved IACUC protocols at the University of Michigan.

### Perfusion of vasculature

To perfuse blood, mice were euthanized then washed with 70% ethanol. The thoracic cavity was opened through the diaphragm, and ribs were cut bilaterally to open the anterior ribcage, which was held with a hemostat. A butterfly needle was injected into the left ventricle and secured. A small incision was made in the right atrium to allow blood to escape, and the mouse was positioned to facilitate drainage from the thoracic cavity. The butterfly needle was connected to a perfusion pump filled with sterile PBS, which was perfused at a rate of 200 mL/h for 15 minutes.

### Tissue harvest and digestion

Synovium was dissected from the medial, lateral, and anterior compartment of the knee, including the fat pad, as described previously^20, 21^. Synovia were digested for 35 minutes in a 1.5 mL mixture containing DMEM with 400 μg/mL collagenase IV, 400 μg/mL liberase, 400 μg/mL DNaseI. 10 seconds of vortexing was performed at 0, 15 and 30 minutes.

Whole joints were isolated and digested in two stages, according to a detailed, published protocol from Leale *et al*^22^. Hindlimbs from the mouse were removed by cutting the proximal leg muscles and dislocating at the hip. Muscles from around the knee joint were trimmed, taking care not to disturb the joint capsule, using a dissection microscope. PBS was used to keep the tissues moist throughout dissection. The knee joint was dislocated at the femoral and tibial growth plates to obtain just the whole joint, which was then transferred to a petri dish for mincing with a scalpel. Minced joint tissue was subjected to 30 minutes of digestion, shaking at 90 rpm at 37°C. This first digestion, to yield the soft tissue fraction, used a 5 mL mixture comprised of 1% collagenase IV (w/v) and 400 µg/mL DNaseI in DMEM containing 5% fetal calf serum. After 30 minutes, the digestate was passed over a 70 µm strainer to collect undigested tissue. Undigested tissue was rinsed with DMEM and 5% fetal calf serum, then digested for a further 90 minutes at 90 rpm and 37°C. This second digestion, to yield the hard tissue fraction, used a 5 mL mixture comprised of 2% collagenase II (w/v) and 400 µg/mL DNaseI in DMEM with 5% fetal calf serum. At the conclusion of soft and hard tissue digests, single cell suspensions were washed in DMEM containing 5% fetal calf serum then subjected to red blood cell lysis with cold ACK lysis buffer before proceeding to flow cytometry staining.

### Flow cytometry

After digestion of synovium or whole joints, single cell suspensions were washed with FACS buffer (PBS containing 2% fetal calf serum and 1 mM EDTA) and 1 µL of FcX TruStain PLUS (Biolegend) was added to each sample. Cells were stained for 30 minutes at 4°C in the dark using fluorescently conjugated antibodies (Supplementary Table 1). After staining, cells were washed with FACS buffer and passed through 35 µm strainers into tubes for flow cytometry. TOPRO3 dye was added for determination of viability. An unstained control was included for all experiments, as well as single-stained control tubes and fluorescence-minus-one tubes for setting negative and positive populations for each color. Flow cytometry was performed on a BD LSRFortessa machine using FACSDiva software for data acquisition.

To visualize high-parameter flow cytometry data in two dimensions, dimensionality reduction was performed using the t-distributed Stochastic Neighbor Embedding (t-SNE) algorithm in FlowJo v10^23^. The automatic learning configuration was used (opt-SNE) and parameters were set to: iterations 1000; perplexity 30; learning rate 5498; KNN algorithm, Exact (vantage point tree); gradient algorithm, Barnes-Hut. All analysis was performed in FlowJo v10 (BD/TreeStar).

### Single-cell analysis of synovial immune cells

Data for the scRNA-seq analysis of synovial immune cells in the ACLR model of PTOA was drawn from GSE211584. Two biological replicates, each comprised of a male and a female synovium, were present for each of the following conditions: Sham (healthy, uninjured), 7d ACLR, and 28d ACLR. Quality control and filtering were performed using Seurat (R, v4.1.0) as described in the original publication^21^. As shown in Figure S1A, immune cell clusters (2, 5 and 8) were computationally subset from all synovial cells. Re-clustering and non-linear dimensionality reduction using uniform manifold approximation and projection (UMAP) was undertaken with dims = 1:35 and res = 0.2. Cluster identities were designated based on the expression of marker genes using the *FindAllMarkers* function and using the Cluster Identity PRedictor tool (CIPR) ^24^. Due to the low abundance of certain immune cell types and difficulty resolving between subsets of cell types, the *CellSelector* tool in Seurat was used to designate cluster identities in a supervised manner based on known marker genes.

### Recapitulation of publicly available datasets

We searched the NCBI Gene Expression Omnibus database using the search terms ‘arthritis’ and ‘synovium’. Datasets from *mus musculus* that did not exclude hematopoietic cells were retrieved and read into RStudio using Seurat. Quality control and filtering were undertaken faithful to the parameters and details provided in the originating publication methods. Where parameters were not provided, defaults were used: *min.cells* = 200, *min.features* = 200, *max.features* = 90^th^ percentile cutoff, *mito.ratio* < 10%. Counts were normalized with the *NormalizeData* function using the *LogNormalize* method, then integrated using RPCA with *variable features* = 3000. Recapitulations for each dataset can be found in Figures S2 (Culemann *et al*), S3 (Sebastian *et al*) and S4 (Muench *et al*).

GSE134420, a scRNA-seq dataset produced by Culemann *et al*^25^, was comprised of sorted CD45+ CD11b+ Ly6G-cells (macrophages/monocytes) from hindpaw synovia of mice subjected to the K/BxN serum transfer arthritis model of RA. No parameters were provided for quality control, so the default parameters above were used. For non-linear dimensionality reduction with UMAP, seven clusters emerged (dims = 1:30, res = 0.15). One cluster was identified as stromal cells (cluster 6), so was removed. Remaining clusters (immune only: 0, 1, 2, 3, 4, 5) were computationally subset and re-clustered using dims = 1:30 and res = 0.15, yielding five clusters, each with unique markers broadly corresponding to the original publication, although the Stmn1+ and Acp5+ macrophages fell into the same cluster.

GSE200843, a scRNA-seq dataset produced by Sebastian *et al*^26^, was comprised of sorted CD45+ cells (all hematopoietic) from whole knee joints of mice subjected to the ACLR model of PTOA. Data were filtered using min.features > 500 and min.cells = 5, in accord with the original publication. We further excluded the top 10% of cells with the highest nFeatures using nFeatures < 4500 and percent.mt < 10, then normalized data using the *LogNormalize* method, and integrated data across all samples using reciprocal PCA, with variable features = 3000. Non-linear dimensionality reduction utilized dims = 1:30 and res = 0.04 to yield seven clusters. A stromal cluster (cluster 4) and an erythrocyte cluster (cluster 6) were detected in the ensuing object, based on marker gene expression. All other clusters (immune only: 0, 1, 2, 3, 5) were then computationally subset and re-clustered using dims = 1:25 and res = 0.07 to yield seven clusters.

GSE184609, a scRNA-seq dataset produced by Muench *et al*^27^, was comprised of sorted live cells from hindpaw synovia of mice subjected to the GPI-induced model of RA. The authors originally used dims = 1:20 for non-linear dimensionality reduction, and filtered based on nFeature and percent.mt cutoffs; however, the values for cutoffs and resolution were not provided. To recapitulate their data most faithfully, we used dims = 1:30 and res = 0.4, and obtained 21 clusters, representing all live sorted cells from the hindpaw synovium. Based on marker gene expression and cluster identity prediction by CIPR, we excluded non-immune cell clusters (1, 3, 10, 11, 12, 13, 14, 18, 19, 20) and computationally subset only the immune cells (0, 2, 4, 5, 6, 7, 8, 9, 15, 16, 17). Immune cells were re-clustered using dims = 1:30 and res = 0.15, yielding eight clusters.

For detailed experimental methods, the reader is directed to the original cited publications for each dataset.

### Integration of datasets

The recapitulated immune objects for each dataset above were integrated using reciprocal PCA in Seurat with variable features = 3000. Non-linear dimensionality reduction was performed using dims = 1:30 and res = 0.2 to generate an integrated UMAP plot for all datasets. The resulting ten clusters were annotated using *FindAllMarkers* and CIPR. Mast cells, due to their very low abundance, were clustered separately in a supervised manner using *CellSelector*, based on expression of marker genes *Il4*, *Kit*, *Fcer1a* and *Mcpt4*.

### Analysis of macrophages in OA and RA datasets

To compare macrophages and monocytes between disease states, two clusters were computationally subset from the integrated object for all datasets (the macrophage and monocyte clusters). Non-linear dimensionality reduction was used to generate a UMAP with dims = 1:20 and res = 0.05, containing three clusters that were annotated using *FindAllMarkers* output. To assess differentially expressed genes (DEGs) in the macrophage cluster between RA (Culemann and Muench) and PTOA (Sebastian and Knights), *FindMarkers* was used. First, macrophages from control vs RA disease, and from control vs PTOA disease were compared to yield DEGs. Using a cutoff of padj < 0.05, the DEGs from both control vs disease comparisons were then compared for overlapping genes and their directionality. To assess differentially regulated pathways in RA or PTOA macrophages, we used PantherDB^28^ and the Gene Ontology database. First, DEGs unique to the RA vs control comparison, and unique to the PTOA vs control comparison, were separately entered into PantherDB for statistical overrepresentation analysis (Gene Ontology: Biological Pathways). To assess conserved pathways in RA and PTOA macrophages, we took DEGs common to both comparisons and with the same directionality and performed statistical overrepresentation testing using GO: Biological Pathways and GO: Molecular Function on PantherDB. All pathway analyses were performed using Fisher’s Exact testing and False Discovery Rate (FDR) was calculated to generate adjusted *P* values (padj). Data were expressed as bubble plots using *ggplot2* v3.4.2^29^.

### Macrophage clustering and trajectory analysis

Myeloid cell clusters from our scRNA-seq data were computationally subset from all immune cells in Seurat, then were exported into Monocle3^30^ for re-clustering to uncover subsets and perform trajectory analysis. Mast cells, DCs and granulocytes were removed from the myeloid object, leaving only monocytes and macrophages, to mitigate the confounding variance from disparate cell types. The monocytes and macrophages were clustered with 15 dimensions, and *cluster_cells()* parameters of resolution = 2.5e-4 and k = 11. For differentiation trajectory analysis, default *learn_graph* arguments were applied to generate a trajectory trail map, which robustly sequestered into two partitions. Differential gene expression analysis and functional pathway analysis were then performed on the three clusters confined to the first partition, containing cells with hallmarks of resident macrophages. Pairwise differential expression analyses were performed with the basal resident, resident-like A, and resident-like B macrophages clusters using *FindMarkers* in Seurat. To assess enriched pathways in each of these three clusters, DEGs from the prior pairwise analyses were used for statistical overrepresentation testing in PantherDB. Biological pathways enriched in any given cluster are indicated in figure panels, derived from genes that were up-regulated in that particular cluster compared to the other. Pathway analyses were performed as described above in comparisons of PTOA and RA macrophages.

For more detailed analysis of how monocytes differentiate into infiltrating macrophages in synovium, we performed gene module analysis. The partition containing the monocytes and infiltrating macrophages was subset using *choose_graph_segments()* and reclustered using 13 dimensions and a resolution of 1e-3 in *cluster_cells()*. An ncenter of 300 and minimal branch length of 15 were applied to generate trajectories. Gene modules, which are groups of co-regulated genes across a pseudotime trajectory, were identified using a resolution of 7e-3. Pseudotime regression plots of key genes were created using the *plot_genes_in_pseudotime* function.

### Transcription factor binding motif analysis

Genes from the three modules with high expression exclusively in the infiltrating macrophage cluster (the terminus of the monocyte-to-infiltrating macrophage trajectory) were submitted to RcisTarget^31^ for transcription factor binding motif analysis. The mm9-tss-centered-5kb-7species.mc9nr.genes_vs_motifs.rankings.feather dataset was used to infer TFs responsible for upregulation of those genes. Significant, direct annotation-derived, high-confidence TF motifs (NES > 3) expressed by at least 5% of macrophages and monocytes with a logcount > 2 were studied. Select gene module genes with a maxRank of 2000 in these motifs were plotted into a network diagram using the visNetwork package.

### Intercellular communication analysis

CellChat^32^ was used to infer ligand-receptor communications between stromal and immune cells clusters. In the *computeCommunProb()* function, the triMean-type analysis was applied. A minimum cell threshold of 10 was applied for each cluster and a significance threshold of 0.05 was applied when inferring communicating pathways. Line plots, heatmaps, and river plots were generated for each condition (Sham, 7d ACLR, and 28d ACLR). Significantly communicating pathways were ranked by ratio of crosstalk communication probabilities (stromal to immune or immune to stromal) relative to all communication probabilities to determine pathways most relevant to crosstalk. Highly ranked pathways were compared between Sham and ACLR conditions to identify crosstalk-relevant pathways most activated by injury. Hierarchy plots of significant communications were generated for CSF, which was identified as a top crosstalk pathway.

CellChat was used to infer outgoing signaling from macrophages, monocytes, and osteoclasts in OA (Knights, Sebastian) or RA (Meunch, Culemann). The triMean type was used in the *computeCommunProb()* function, with a minimal cell count of 10 per cluster. Significant pathways (*P* < 0.05) were included for line plot, heatmap, and river plot generation. A default cutoff of 0.5 was used for river plots of significant outgoing communications.

## RESULTS

### Diverse immune cell types are present in healthy and injured synovium

Having characterized the stromal niche of synovium in PTOA and defined distinct functional subsets of fibroblasts^21^, we sought to understand the immune profile of healthy and injured synovium. Using our published scRNA-seq data of synovial cells from Sham, 7d ACLR or 28d ACLR mice (GSE211584), we computationally subset only immune cells based on their cluster identity and verified their positivity for *Ptprc* expression (encoding CD45) and negativity for *Pdgfra* expression (encoding PDGFRα) (Fig S1A-B). In the absence of stromal and vascular cells that would otherwise confound variance, re-clustering was performed on the immune cells, revealing a diverse profile of innate and adaptive cell types, including monocytes and macrophages, dendritic cells (DCs), T cells, B cells, mast cells, granulocytes, innate lymphoid cells (ILCs) and natural killer (NK) cells (Fig 1A, Fig S1C). Abundance of synovial immune cells greatly increased acutely after injury (7d ACLR), with a degree of resolution occurring by 28d ACLR (Fig 1B-C). Macrophages, unsurprisingly, predominated both numerically and proportionally in healthy and injured synovium, with various subsets of T cells and DCs comprising most of the remaining cells (Fig 1C, Fig S1D-F). To assess whether the immune cell types observed in scRNA-seq based on their transcriptomic signatures were detectable at the functional protein level using surface markers, we employed multi-color flow cytometry. Indeed, all major groups of immune cells were identified within the CD45+ pool of synovial cells, based on their surface marker and light scatter profiles (Fig 1D-E).

**Figure 1.**
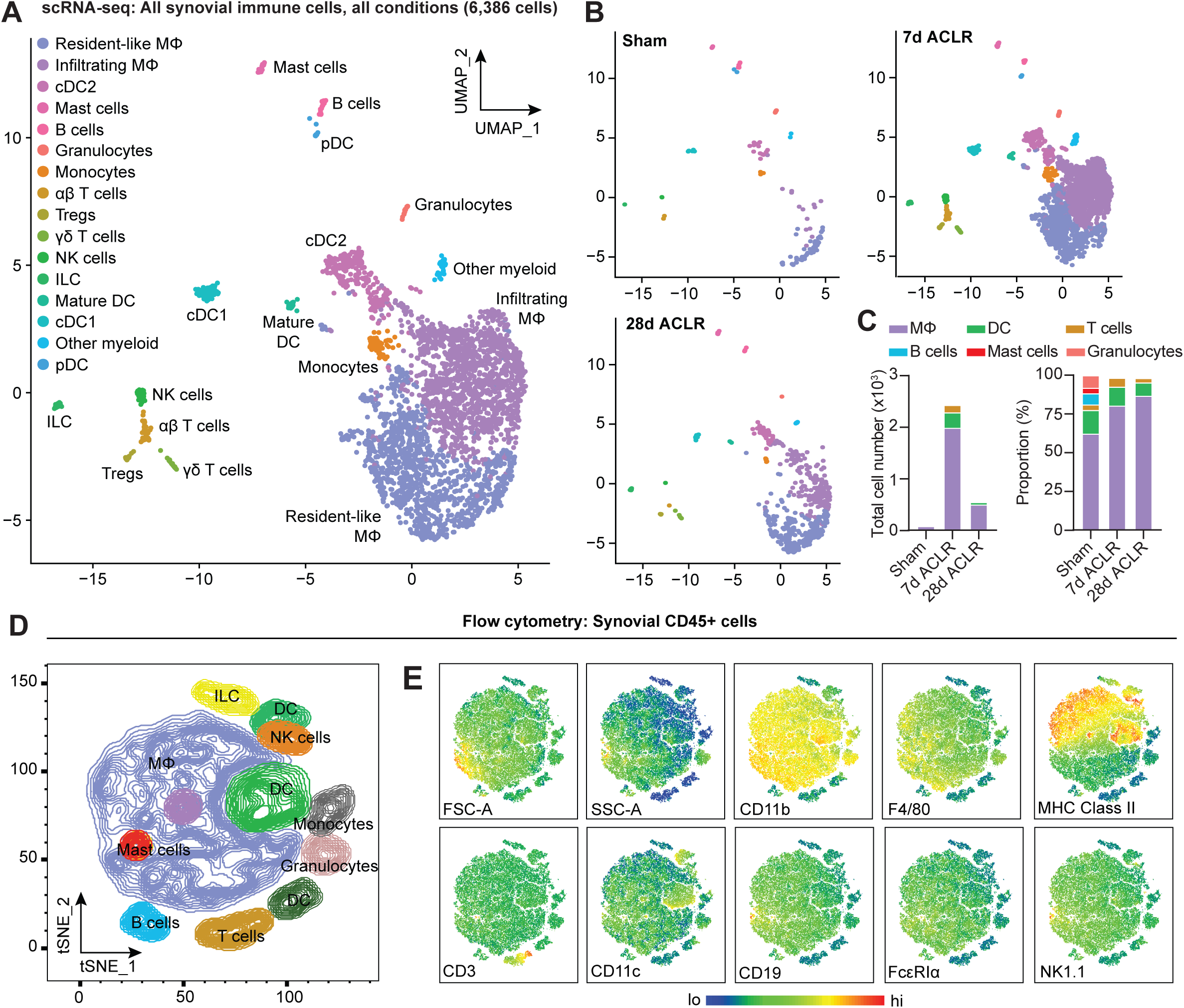
Diverse immune cell types are present in healthy and injured synovium. (A) UMAP plot of all immune cells by scRNA-seq of synovium from mice subjected to Sham, 7d ACLR and 28d ACLR, or (B) split by condition. (C) Breakdown of immune cell types by total abundance (left) or proportion (right) in Sham, 7d ACLR or 28d ACLR synovium. (D) t-SNE plot of immune cell types by flow cytometry of CD45+ synovial cells from mice subjected to Sham (left and right synovia from n=4 mice) and 7d ACLR (right synovia from n=4 mice). (E) t-SNE heatmaps of scatter and surface marker parameters used to define synovial immune cell identities by flow cytometry. Tregs: regulatory T cells; NK cells: natural killer cells; ILC: innate lymphoid cells; DC: dendritic cells; cDC: conventional DCs; MΦ: macrophages; FSC-A: forward scatter-area; SSC-A: side scatter-area.

### Integration of joint immune cells from mouse scRNA-seq datasets

Given the recent proliferation of studies seeking to describe the immune landscape in murine arthritis models, there is a strong rationale for harmonizing this publicly available data to characterize similarities and differences. We obtained four published scRNA-seq datasets for integration and further analysis of the immune cell niche: Culemann *et al*^25^ (GSE134420; sorted CD45+ CD11b+ Ly6G-cells from hindpaw synovia of mice subjected to the K/BxN serum transfer arthritis model of inflammatory arthritis/RA); Sebastian *et al*^26^ (GSE200843; sorted CD45+ cells from whole knee joints of mice subjected to the ACLR model of PTOA); our data, from Knights *et al*^21^ (GSE211584; knee synovium from mice subjected to the ACLR model of PTOA); and Muench *et al*^27^ (GSE184609; sorted live cells from hindpaw synovia of mice subjected to the GPI-induced model of inflammatory arthritis/RA).

Datasets were obtained from the NCBI Gene Expression Omnibus then recapitulated in accordance with parameters provided in the Methods sections of each paper, including computational removal of any non-immune cell clusters (Fig S2-S4). The resulting immune cell-only objects were integrated using reciprocal PCA and assigned identities based on known marker genes and alignment to published transcriptomic signatures of immune cell types using the Cluster Identity PRedictor (CIPR) ^24^ (Fig 2A-B, Fig S5A). Monocytes, macrophages, DCs, neutrophils, B cells, T and NK cells, mast cells, and proliferating immune cells were all detected, and the same, or very similar, cell types clustered closely together across datasets following integration (Fig 2A-D). Likely owing to the disparate models (inflammatory arthritis/RA or PTOA), anatomical locations (knee synovium, whole knee joint, or hindpaw synovium), and experimental approaches (eg. sorting only monocytes and macrophages), certain cell types predominated in some datasets but were largely absent in others. This was most strikingly evident with the neutrophil clusters (Neut-1, -2 and -3) present primarily in the Sebastian and Muench datasets, the latter of which modeled rheumatoid arthritis in which neutrophils are known to play a larger role^33^. One explanation for high neutrophil abundance in these datasets may be bone marrow contamination, given that these studies digested the whole knee joint (Sebastian) or the whole hindpaw (Muench). We detected a high abundance of neutrophils in digested whole knee joints by flow cytometry - approximately half of all CD45+ cells (Fig S6A). This number was not reduced when mice were systemically perfused to clear the vasculature, suggesting that neutrophils in the joint digest are not derived from local blood vessels. Comparison of the gene signatures of neutrophil clusters 1, 2 and 3 to publicly available neutrophil datasets support that these cells, particularly Neut-1, are more likely bone marrow-derived than true intra-articular tissue neutrophils (Fig S6B-C, Supplementary Table 2).

**Figure 2.**
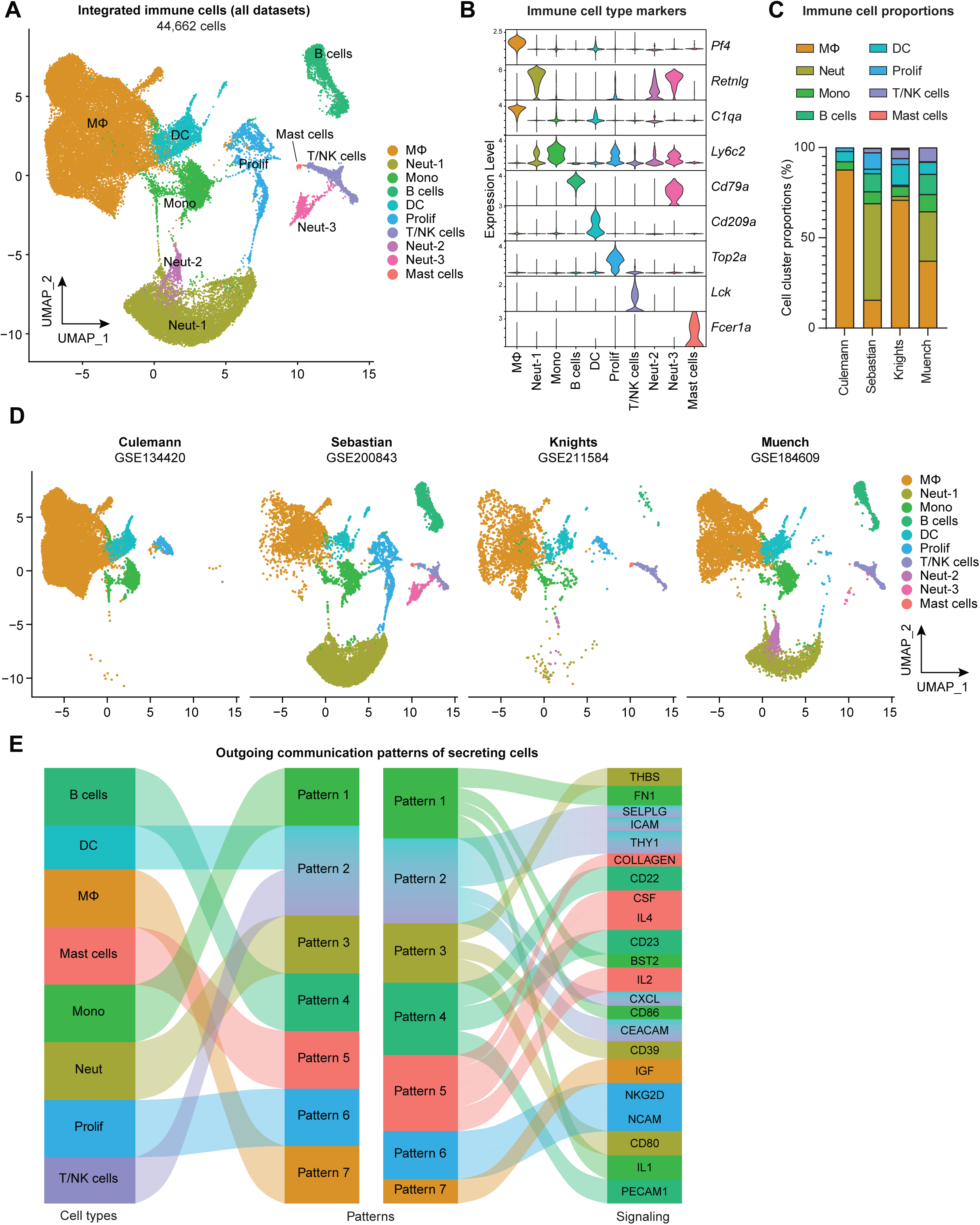
Integration of joint immune cells from mouse arthritis scRNA-seq datasets. (A) Integrated UMAP plot of all immune cells from GSE134420 (Culemann), GSE200843 (Sebastian), GSE211584 (Knights) and GSE184609 (Muench). (B) Violin plots showing gene markers for each cell cluster. (C) Proportional breakdown of each major immune cell type in each dataset. (D) UMAP plots showing immune cell clusters for each dataset. (E) CellChat outgoing communication patterns for each major immune cell group Neut: neutrophils; Mono: monocytes; Prolif: proliferating cells.

Across datasets, myeloid cells (monocytes, macrophages, granulocytes, dendritic cells) were the predominant cell group, with lymphocytes (T and B cells) comprising less than 20% of total immune cells, and mast cells were very rare (Fig 2C, Fig S5B). To assess broad signaling patterns of immune cell types across datasets and disease states, we used CellChat^32^. River plots of outgoing communication signals from immune cell clusters showed distinct broad signaling patterns for most immune cell types across datasets, with the notable exception of DCs and T/NK cells which were all predicted to participate strongly in signaling via integrins, selectins and chemokine ligands (Fig 2E, Fig S5C-D). Mast cells were enriched for sending out CSF and interleukin-4 (IL-4) signals, while monocytes and macrophages were enriched for IL-1 and IGF signaling.

### Comparison of macrophages between disease states

Much attention has been devoted to the similarities and differences between OA and RA^34-37^, however less focus has been directed at how intra-articular macrophages compare between the two diseases. From the integrated object containing immune cell clusters from all four datasets, we subset only monocytes and macrophages (Fig 3A). The resulting three clusters corresponded to monocytes (*Plac8* and *Ly6c2*), macrophages (*C1qa* and *Adgre1*), and osteoclast-like cells (*Ctsk* and *Acp5*) (Fig 3B). Macrophages represented a higher proportion of cells in the inflammatory/RA datasets (Culemann and Muench), while monocytes were more proportionally abundant in the PTOA datasets (Sebastian and Knights) (Fig 3C-D, Fig S7A). Osteoclasts represented <5% of cells, and macrophages were vastly more abundant than monocytes or osteoclast-like cells in both PTOA and RA models.

**Figure 3.**
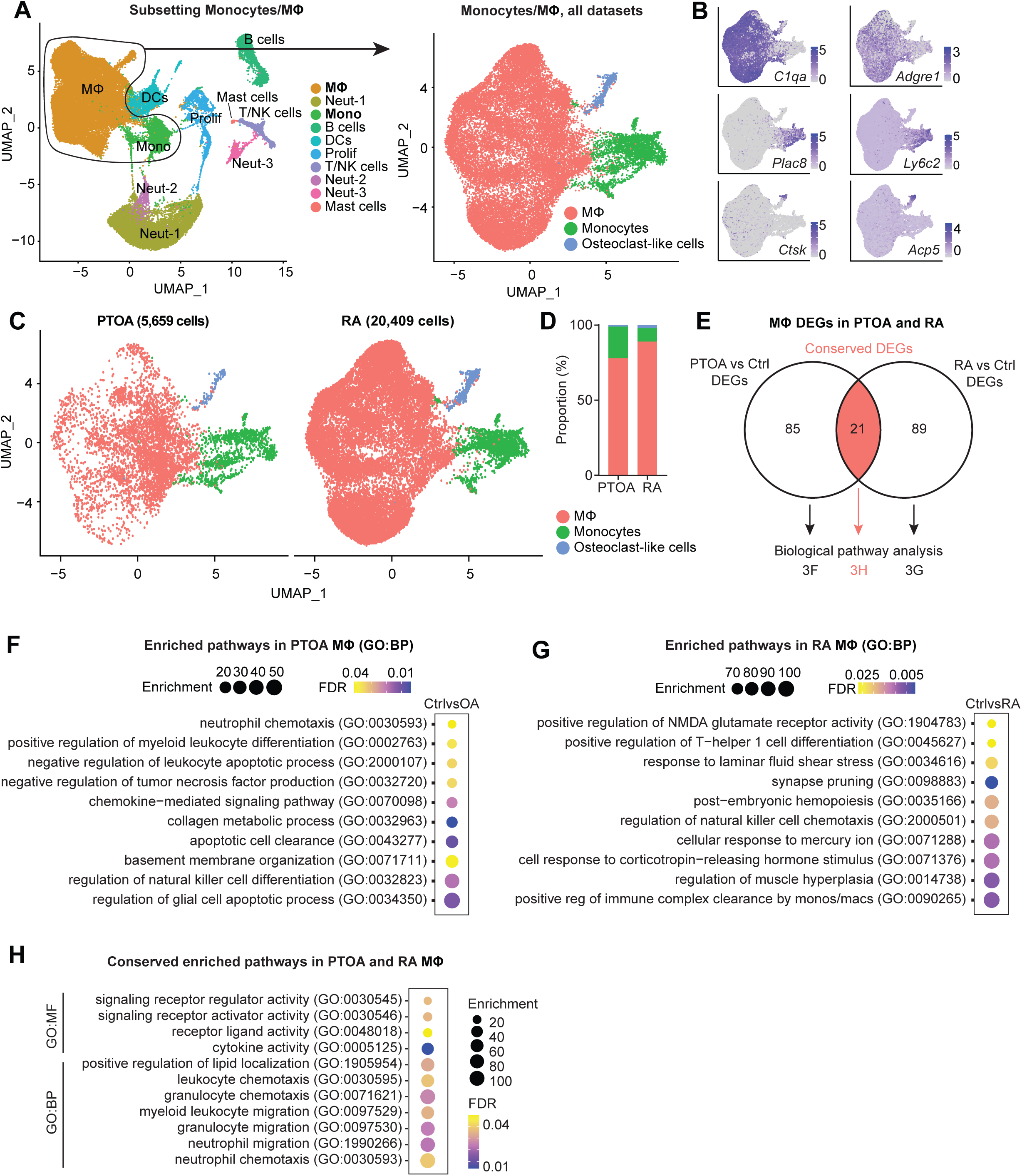
Comparison of macrophages between arthritis disease states. (A) Monocytes and MΦ from all PTOA and RA immune cell datasets were computationally isolated, resulting in three clusters: MΦ, monocytes and osteoclast-like cells. (B) Feature plots showing expression levels of highly enriched genes unique to each cluster. (C) Side-by-side UMAP plot of monocytes and MΦ from PTOA and RA datasets and (D) the proportion of each cluster within each disease state. (E) Differential gene expression analyses were performed on the MΦ cluster specifically. Comparison groups were PTOA vs control (Ctrl) MΦ and RA vs control (Ctrl) MΦ, with overlapping and non-overlapping DEGs for each comparison shown in a Venn Diagram. Also see Supplementary Tables 3 and 4. (F-G) Enriched biological pathways in PTOA (F) and RA (G) MΦ when compared to their respective control MΦ. (H) Conserved enriched biological pathways between PTOA and RA MΦ, derived from common DEGs with the same directionality between both separate comparisons. Also see Supplementary Table 5. For all pathway analyses, statistical overrepresentation tests were performed with Fisher’s Exact testing and calculation of false discovery rate (FDR). GO:BP: Gene Ontology Biological Pathways; GO:MF: Gene Ontology Molecular Function.

CellChat analysis of outgoing signaling patterns for each cluster in PTOA and RA highlighted monocytes as dominant drivers of signaling with a high degree of commonality between diseases (Fig S8). Macrophages and osteoclasts, on the other hand, exhibited more disparate outgoing signaling programs between PTOA and RA. To better understand transcriptomic similarities and differences between macrophages from PTOA and RA synovial joints, we next performed two differential gene expression analyses: i) PTOA versus healthy control macrophages (Sebastian and Knights) and ii) RA versus healthy control macrophages (Culemann and Muench). 85 differentially expressed genes (DEGs) were found exclusively between PTOA and control macrophages, 89 DEGs were found exclusively in the RA versus control comparison, and 21 DEGs overlapped between both analyses (Fig 3E, Supplementary Tables 3 and 4). Thus, to determine perturbed pathways unique to macrophages in each disease state separately, we used exclusive DEGs to perform pathway analysis using Gene Ontology (Supplementary Table 5). In PTOA macrophages, compared to controls, collagen metabolism, myeloid differentiation regulation, neutrophil chemotaxis and glial cell apoptosis pathways were all enriched (Fig 3F). In RA macrophages on the other hand, response to corticotropin-releasing hormone (known to be pro-inflammatory and enriched in RA synovial fluid^38^), Th1 differentiation, and cellular response to mercury were all perturbed (Fig 3G). Analysis of enriched pathways common between RA and PTOA macrophages yielded unsurprising results, including cytokine activity, white blood cell chemotaxis and migration, and signaling receptor activity (Fig 3H).

### Synovial macrophage subsets and trajectories in PTOA

In Figure 1 we identified macrophages and monocytes as the most abundant immune cell group in both healthy and PTOA synovium. To eliminate variation from other cell types and to more sensitively describe transcriptional variance amongst macrophages, we computationally isolated monocytes and macrophages from all immune cells in our scRNA-seq dataset (Fig 4A, Fig S9A). Five clusters emerged, expressing classical monocyte and macrophage markers *Itgam* (encoding CD11b), *Adgre1* (encoding F4/80), *Csf1r* (encoding the M-CSF receptor) and *Cx3cr1* (Fig 4A, Fig S9B). Sham synovium was predominated by macrophages expressing *Timd4*, *Lyve1* and *Folr2* which we have termed basal resident macrophages, given their resemblance to homeostatic macrophages residing in various tissues with the same gene expression signature^39, 40^ (Fig 4B-C). After injury, there was a profound increase in cellular abundance and heterogeneity, but with the notable loss of cells characterized by the basal resident gene signature. Trajectory modeling using Monocle3 indicated that basal resident macrophages polarized into distinct phenotypes in PTOA, characterized by expression of *Mrc1*, *Ly6e* and *Gas6* (resident-like A) or *Cd9*, *Spp1* and *Trem2* (resident-like B) (Fig 4C-D).

**Figure 4.**
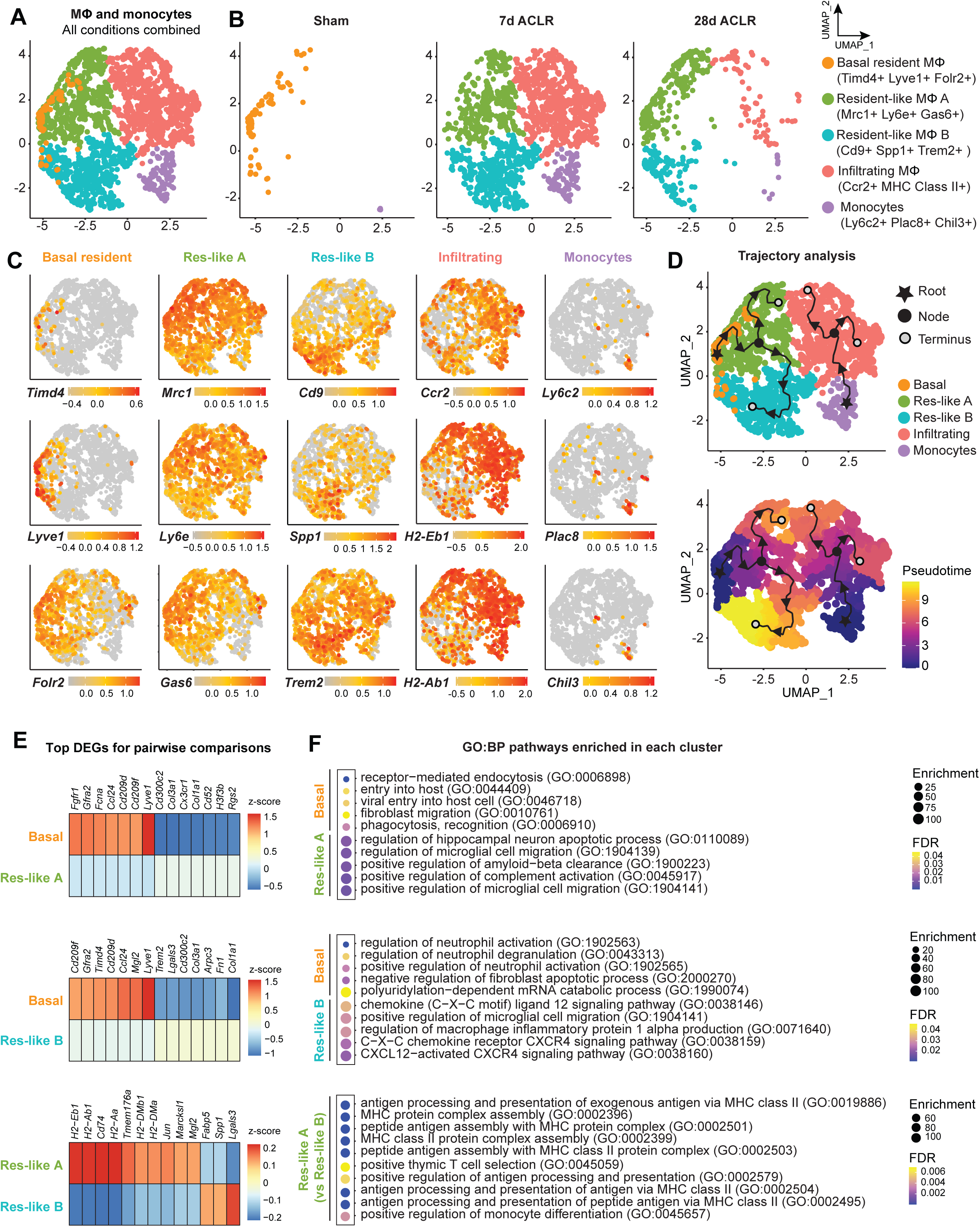
Synovial macrophage subsets and trajectories in PTOA. (A) UMAP plot of monocytes and MΦ from synovium of Sham, 7d ACLR and 28d ACLR mice, or split by condition (B). Cluster naming and top gene markers are given on the right. (C) Gene feature plots showing expression of key marker genes for each subset. (D) Pseudotime trajectories overlaid onto monocyte and MΦ subsets showing directionality (arrowheads), starting points (roots, stars), branching points (nodes, black circles), and endpoints (termini, grey circles with black outline). Partitions are shown as disconnected (separate) trajectory trails. Colored cell clusters are shown in the top plot and pseudotime scale is shown in the bottom plot. (E) Heatmaps of top differentially expressed genes (DEGs) in pairwise comparisons for basal resident MΦ, resident-like MΦ A, and resident-like MΦ B clusters (Padj < 0.05). (F) Enriched biological pathways in basal resident MΦ, resident-like MΦ A, and resident-like MΦ B clusters, derived from statistical overrepresentation tests of DEGs from corresponding pairwise comparisons in (E). Also see Supplementary Table 6. Fisher’s Exact testing was performed and false discovery rate (FDR) was calculated. GO:BP: Gene Ontology Biological Pathways.

The top DEGs between resident-like macrophage clusters were analyzed to better understand their potential functional differences (Fig 4E, Supplementary Table 6). Compared to basal resident macrophages, the resident-like A cluster was enriched for functions pertaining to complement activation and cell migration (Fig 4F). Resident-like B cells had enriched CXCL12-CXCR4 signaling and production of macrophage inflammatory protein, compared to the basal resident cells. On the other hand, compared to the two polarized resident-like macrophage subsets, basal resident cells had functional enrichment of homeostatic biological pathways including phagocytosis, endocytosis, and apoptosis. The resident-like A subset was enriched for antigen presentation functions compared to resident-like B.

Monocytes expressing *Ly6c2*, *Plac8* and *Chil3* were also present, particularly after injury, and trajectory analysis suggested that they give rise to a pro-inflammatory cluster expressing *Ccr2* and genes encoding MHC Class II, which we have termed infiltrating macrophages. Genes historically used to mark ‘M1’ or ‘M2’ macrophages, including *Tnf* (encoding TNF-α) and *Il10* (encoding IL-10), did not segregate neatly into defined clusters (Fig S9C-D), emphasizing the need to supersede these outdated terms with more nuanced characterizations based on their *in vivo* gene expression and functional profiles.

Together these results suggest that functional polarization states of macrophages arise from a resident pool of cells during PTOA, and that a pro-inflammatory infiltrating macrophage population derives from circulating monocytes. Importantly however, *in vivo* approaches such as lineage tracing will be required to assess the veracity of *in silico* trajectory modeling.

### Stromal-immune crosstalk via M-CSF signaling

Stromal and immune cells participate in autocrine and paracrine signaling within synovium, and we sought to define key signaling axes that regulate the synovial immune cell niche. Using CellChat ligand-receptor interactions for all synovial cell types defined as stromal or immune, we implemented a novel approach to derive crosstalk communication scores for multi-directional and uni-directional stromal-immune signaling (Fig 5A). The M-CSF pathway, involving the ligands M-CSF (macrophage colony stimulating factor) and IL-34, and the M-CSF receptor, was highly induced after injury, but primarily in the stromal-to-immune direction (Fig 5B, Fig S10), implying that stromal-derived ligands regulate synovial immune cells expressing the M-CSF receptor – namely, macrophages. M-CSF and its receptor are well-recognized for their role in monocyte and macrophage function^41, 42^, and more recently, IL-34 was identified as an alternative M-CSF ligand^43, 44^. Bulk RNA-sequencing of synovium showed robust induction of *Csf1* (encoding M-CSF), *Il34* (encoding IL-34) and *Csf1r* (encoding the M-CSF receptor) after joint injury (Fig 5C); thus, we sought to determine the cell types orchestrating this signaling axis.

**Figure 5.**
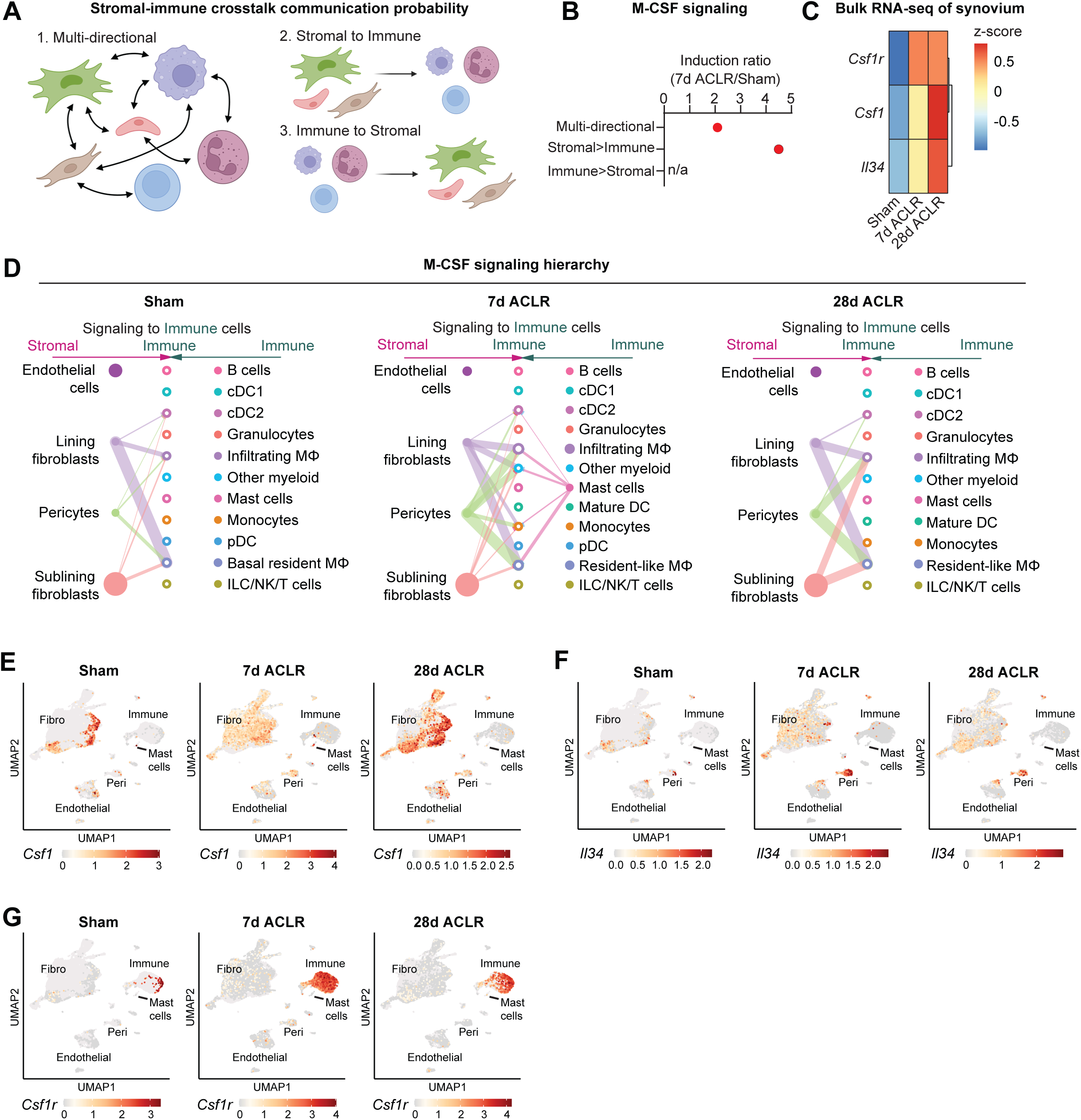
Stromal-immune crosstalk via M-CSF signaling. (A) Cartoon depicting multi-directional and uni-directional crosstalk between stromal and immune cells in synovium, used for calculation of crosstalk communication probability scores. (B) Induction ratio for multi-directional and uni-directional M-CSF signaling derived from crosstalk communication probability scores (7d ACLR/Sham). (C) Expression of M-CSF ligands *Csf1* and *Il34*, and the M-CSF receptor *Csf1r*, from bulk RNA-seq of Sham, 7d ACLR, or 28d ACLR synovium (n=5-6 male and n=5-6 female synovia per condition) (dataset available at NCBI GEO *–* upload and acceptance pending; accession number will be added upon receipt). (D) CellChat hierarchy plots for the M-CSF signaling pathway in Sham, 7d ACLR and 28d ACLR. Circles with fill represent cells sending signals, circles without fill represent cells receiving signals, in each condition. Line thickness corresponds to strength of communication. (E-G) Feature plots showing expression of *Csf1* (E), *Il34* (F) and *Csf1r* (G) in all synovial cells, split by condition (Sham, 7d ACLR, or 28d ACLR). n/a: not applicable; Fibro: fibroblasts; Peri: pericytes.

CellChat hierarchy plots, which infer directionality and degree of signaling between each cell type based on ligand-receptor interactions of a given pathway, were generated for all synovial cells from Sham, 7d ACLR, and 28d ACLR. In Sham synovium, lining and sublining fibroblasts, as well as pericytes, were the primary producers of M-CSF and/or IL-34 ligand, with macrophages and dendritic cells being the primary signal-receiving cell types (Fig 5D). After injury, alongside the increased cell type diversity, there were more M-CSF pathway interactions (number of connections) with higher probabilities (thickness of connections), which, again, originated from fibroblasts and pericytes and signaled towards myeloid cells. A small population of mast cells also sent outgoing signals at 7d ACLR only. No immune-to-stromal signaling via the M-CSF pathway was predicted (Fig S11). Expression patterns of M-CSF pathway genes in synovial cell types confirmed fibroblasts as the major source of *Csf1* ligand, with no expression seen in immune cells (except for mast cells); pericytes were the primary source of *Il34* ligand; and *Csf1r* expression was confined to the immune cell compartment (Fig 5E-G). These results shed light on the cellular participants, temporal dynamics, and regulation of M-CSF signaling in synovium, which is among the most activated stromal-immune crosstalk axes following joint injury.

### Transcriptional control of monocyte differentiation in synovium

Canonical monocyte to macrophage differentiation is well characterized, however the regulation of this phenomenon in synovium remains largely undescribed. Thus, we next sought to model the differentiation mechanism of blood-derived monocytes that enter synovium and give rise to infiltrating macrophages. The trajectory from Figure 4D was isolated to only include monocytes to infiltrating macrophages, eliminating unrelated variance stemming from resident macrophage clusters (Fig 6A-B). Modules of genes co-regulated across pseudotime were generated using Monocle3 (Fig 6C, Supplementary Table 7), which represent the various transcriptional programs activated during monocyte-to-macrophage maturation, and the modules most highly enriched at the trajectory terminus were further analyzed (modules 1, 4 and 6).

**Figure 6.**
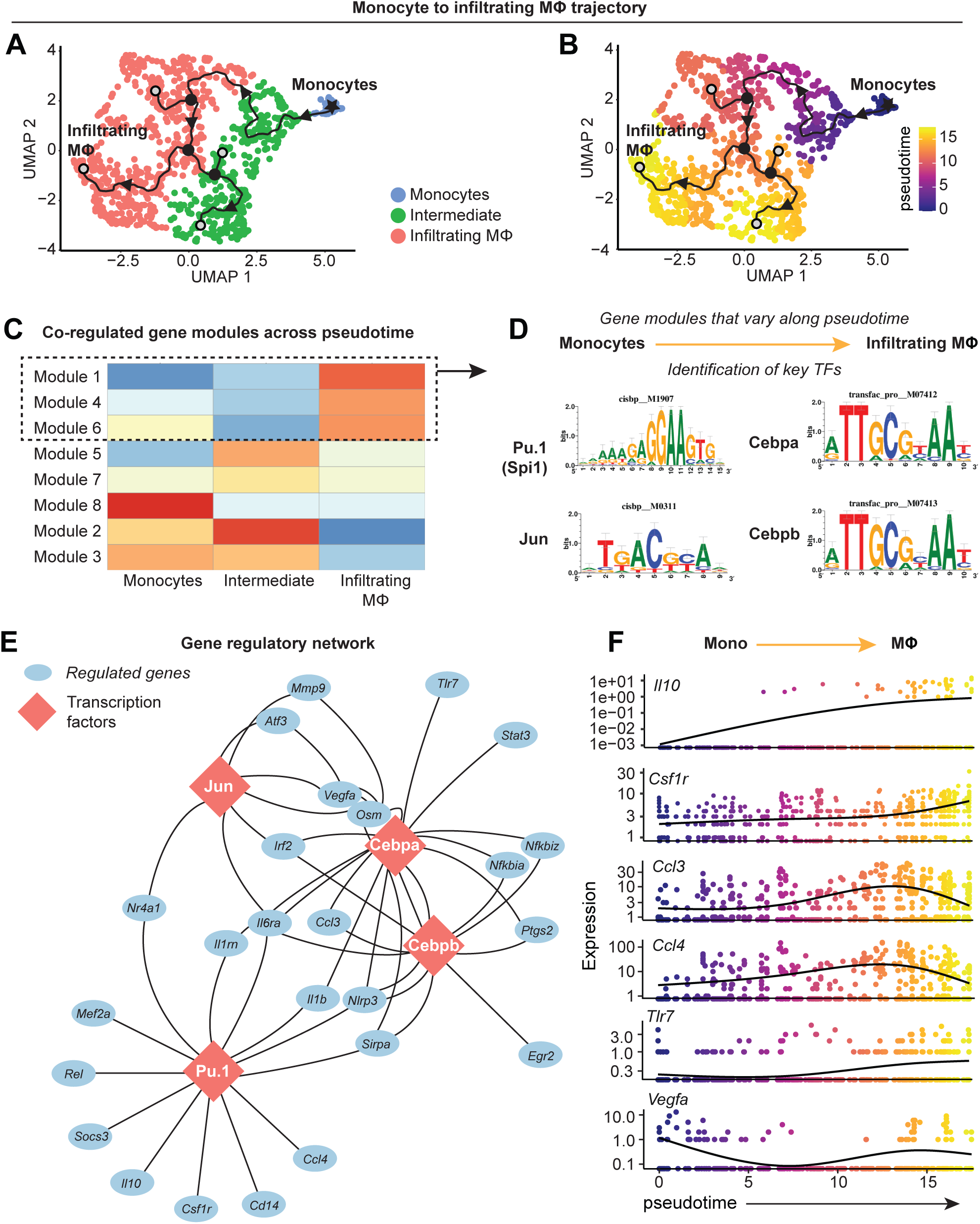
Transcriptional control of monocyte differentiation in synovium. (A-B) Pseudotime trajectory from monocytes to infiltrating MΦ showing directionality (arrowheads), starting point (root, star), branching points (nodes, black circles), and endpoints (termini, grey circles with black outline). Pseudotime scale is shown in (B). (C) Heatmap of gene module analysis of co-regulated genes across pseudotime trajectory from monocytes to infiltrating MΦ. Also see Supplementary Table 7. (D) Modules 1, 4 and 6 were subjected to promoter screening for putative transcriptional regulators of module genes, using RcisTarget. Top directly-annotated motif hits and their corresponding transcription factors (TFs) are shown. (E) Gene regulatory network for the TFs Pu.1 (Spi1), Jun, Cebpa, and Cebpb. (F) Pseudotime regression plots of selected gene from the gene regulatory network in (E). Monocytes to infiltrating MΦ, left to right.

Given that genes in these modules are induced in the terminal stage of monocyte-to-macrophage differentiation in synovium, we screened for which transcriptional factors (TFs) control this fate. Genes from modules 1, 4 and 6 were analyzed by RcisTarget, which screens gene promoters for overrepresented binding motifs, allowing us to infer TFs that may putatively control monocyte-to-macrophage differentiation in synovium. After filtering out putative factors showing little or no expression, the top-ranked candidates for regulating differentiation were Pu.1 (Spi1), Cebpα, Cebpβ, and Jun (Fig 6D), all of which have reported roles in controlling macrophage fate and function^45-48^. The gene regulatory network for these TFs showed distinct (eg. *Il10*, *Rel*, *Tlr7*, *Egr2*) and shared (eg. *Vegfa*, *Irf2*, *Il6ra*, *Nlrp3*) target genes, with Pu.1 controlling expression of *Csf1r* as has been described in other settings^45, 49^. Expression of the genes encoding Pu.1, Cebpα, Cebpβ, and Jun was not necessarily induced over pseudotime, however their gene targets (derived from modules 1, 4 and 6), such as *Il10*, *Csf1r*, and *Tlr7*, showed increasing expression with pseudotime (Fig 6F and Fig S12). Together, these results point towards four key TFs that control the differentiation of blood-derived monocytes into pro-inflammatory synovial macrophages and identified the genes they regulate during this process, while corroborating our understanding of canonical monocyte differentiation and how it is controlled.

## DISCUSSION

Chronic synovitis is recognized to drive various pathological processes associated with arthritis, including pain and tissue damage, but the early cellular and molecular events following joint injury are not well described. We recently demonstrated that synovial fibroblasts expand rapidly and phenotypically diverge into distinct functional subsets in PTOA, alongside pathological activation of canonical Wnt signaling^21^. The present work demonstrates the injury-induced immune cell dynamics concurrent with the activation of synovial fibroblasts, and our flow cytometric and transcriptomic analyses demonstrate that while most immune cells are rare in healthy synovium, joint injury causes massive diversification and influx of immune subsets. Macrophages exhibited the greatest expansion, and our single-cell transcriptomic analyses identified a resident macrophage population that polarizes into two phenotypes following injury, in addition to an infiltrating macrophage population derived from monocytes. Synovial stromal cells were highly active in mediating immune cell infiltration and expansion, with unbiased modeling of stromal-immune crosstalk identifying that M-CSF signaling is among the most perturbed ligand-receptor axes after injury. The TFs Jun, Cebpα, Cebpβ, and Pu.1 were identified as putative mediators of differentiation from monocytes to synovial macrophages.

Among the most pressing questions in the field of PTOA research is which immune cells drive disease versus which promote resolution following joint injury. Rigorously describing immune cell dynamics and the relative contribution of specific cellular subsets to disease processes such as pain, osteophyte formation, and cartilage damage will facilitate the development of targeted, disease-modifying therapies. Consistent with prior studies^26, 50, 51^, our transcriptomic profiling demonstrated that healthy synovia contained all major immune cell types, including rare subsets such as mast cells and T cells, and this was corroborated by flow cytometry. As we did not perfuse mice prior to tissue collection, some of these may be blood-derived, but mature immune cells such as mast cells, T cells, B cells, and dendritic cells are unlikely to be present in healthy synovial tissue vasculature in sufficient numbers to generate this result. Robust lineage tracing studies are lacking to describe the origin, maintenance, and replenishment of these immune cells, which will require more specific tools given that recent scRNA-seq studies have revealed that many of the existing genetic reporter systems are insufficient to distinguish between resident versus systemically-derived cells, or exhibit too much overlap in expression among different immune subsets (e.g. *LysM-cre*, *Cx3cr1-cre*, *Ccr2-cre*).

Macrophages were the most numerous immune cell type in healthy and injured synovium and exhibited the greatest expansion following injury. In healthy tissue, we identified a resident synovial macrophage population with a *Timd4+ Lyve1+ Folr2+* (TLF) signature that resembled homeostatic tissue-resident macrophages described in other tissue contexts^39, 40^. Trajectory analysis predicted that these resident macrophages could expand into two injury-induced resident-like subsets: one marked by *Mrc1* (encoding CD206) and *Gas6*, and another marked by *Cd9, Spp1* and *Trem2* expression. Several recent studies have corroborated the existence of a pro-fibrotic macrophage population that emerges in different disease contexts^4, 11, 52-57^, expressing *Spp1* (encoding osteopontin), *Cd9* and *Trem2*, strongly suggesting that the resident-like B macrophage population we identified in PTOA synovium is functionally analogous, which is significant in light of the strong fibrotic phenotype of PTOA synovium. Our trajectory modeling pointed towards these pro-fibrotic macrophages arising from synovium-resident TLF+ macrophages after injury. In liver cirrhosis, however, trajectory analysis suggested a monocytic origin for *Spp1*+ macrophages. Thus, robust *in vivo* fate mapping studies are critical to elucidate their true origins. The resident-like A cluster that emerged after injury expressed *Mrc1* (CD206), historically associated with an anti-inflammatory and reparative phenotype, and which has been shown to mediate collagen turnover^58, 59^. In addition their potentially beneficial role in regulating extracellular matrix, the resident-like A population also expressed *Gas6*, which is required for efficient efferocytosis of apoptotic cells by synovial macrophages^60^ – an important aspect of inflammatory resolution that goes awry in OA^16^. It is noteworthy that conventional “M1” and “M2” markers, such as *Il1b*, *Tnf*, *Arg1* and *Il10*, revealed no appreciable separation across our clusters, placing further impetus to move beyond the simplistic M1-M2 spectrum of macrophages. Mechanistic and *in vivo* studies are now warranted to describe the precise pathophysiological functions of these subsets in OA, their ontogenies, and their spatial distribution in the various synovial niches.

We further identified a clear monocyte-to-macrophage trajectory, and these infiltrating macrophages highly expressed the pro-inflammatory genes encoding for CCR2, IL-1β, TNF-α, MHC Class II, among others. Huang *et al* recently demonstrated in murine inflammatory arthritis that synovial inflammatory resolution is achieved, in part, by suppressing the *in situ* differentiation of infiltrating monocytes to a F4/80^hi^ MHC Class II^+^ phenotype, which would otherwise promote chronic inflammation^17^. Thus, we conclude that the monocyte-derived macrophages we identified to emerge in synovium following joint injury are pro-inflammatory and analogous to those described by Huang *et al*, and developing an understanding of the molecular mechanisms that promote their recruitment and differentiation following joint injury is critical. We therefore undertook further focused analysis of the transcriptional trajectory spanning monocytes to infiltrating macrophages, and we identified Jun, Cebpα, Cebpβ, and Pu.1 as TFs potentially regulating the gene programs of monocyte-to-macrophage maturation in OA synovium. These have been previously attributed to regulate macrophage maturation in other disease contexts such as rheumatoid arthritis^61, 62^, often in concert with each other^63^, however empirical studies demonstrating whether maturation of systemically-derived monocytes into inflammatory synovial macrophages during OA is dependent on these TFs are needed.

To understand which stromal-derived signals recruit or activate immune cells, we extended the widely-utilized cellular crosstalk toolbox CellChat to identify the ligand-receptor axes most perturbed by joint injury and most active in stromal-to-immune communication, in an unbiased fashion. By deriving probability ratios for all possible multi- and uni-directional stromal-immune signaling axes in healthy and injured joints, M-CSF signaling emerged as a major crosstalk axis activated by injury. M-CSF (and its alternative ligand, IL-34) has been shown to mediate monocyte-macrophage maturation via the M-CSF receptor in other disease contexts^41, 49, 64^, and monoclonal antibody-mediated M-CSF receptor inhibition ameliorated cytokine production in human RA synovial explants and mitigated disease severity in murine collagen-induced arthritis^65^. Our TF binding analysis revealed Pu.1 as a putative regulator of *Csf1r*, in agreement with previous reports^45, 49, 66^. Ongoing studies are seeking to understand whether synovial fibroblast-derived M-CSF activates putative regulators like Pu.1, to promote monocyte maturation towards a pro-inflammatory phenotype. Further work is also needed to describe which upstream pathways promote M-CSF and IL-34 ligand secretion by synovial fibroblasts, pericytes and mast cells. Given the universal dependence of macrophages, both resident and recruited, on M-CSF signaling, dissecting the nuanced mechanistic distinctions such as ligand-receptor combination, ligand bioavailability, sending and receiving cell types, and spatial proximity, will be crucial if targeting M-CSF signaling is to be harnessed therapeutically in OA. In conjunction, accurately defining the temporal dynamics, ontogenies, and pathological versus protective functions of distinct macrophage subtypes in synovium will be integral if these cells are to be targeted for intra-articular treatment.

## Data availability statement

All scRNA-seq datasets are publicly available from the NCBI Gene Expression Omnibus using the following accession numbers: GSE134420, GSE200843, GSE211584 and GSE184609. Bulk RNA-seq data is pending upload to NCBI Gene Expression Omnibus.

## Acknowledgements

We would like to acknowledge the services offered by the University of Michigan Flow Cytometry Core.

## Contributions

AJK: conception and design; collection and assembly of data; analysis and interpretation of data; manuscript writing

ECF: collection and assembly of data; analysis and interpretation of data; manuscript writing

OME: collection and assembly of data; analysis and interpretation of data

MS: collection and assembly of data

CTA: analysis and interpretation of data

TM: conception and design; analysis and interpretation of data; manuscript writing; final approval of the article (TM takes responsibility for the integrity of the work as a whole, from inception to finished article)

## Funding sources

This work was supported by funding from the National Institutes of Health to AJK (K99AR081894) and TM (R01AR080035, R21AR076487) and from the Dr. Ralph and Marian Falk Medical Research Trust (Catalyst Award to TM). TM receives further unrelated research support from the National Institutes of Health (R21AR080502, R21AR082016, UC2AR082186) and the Department of Defense/CDMRP (GRANT13696744). AJK was supported by a Pioneer Postdoctoral Fellowship from the University of Michigan. ECF was supported by the Dr. Ralph and Marian Falk Medical Research Trust and National Institutes of Health (R01AR080035 to TM).

## Competing interests

The authors have no competing interests to declare.

## Notes

### Competing Interest Statement

The authors have declared no competing interest.

