## Supplementary figures and legends for "Synovial macrophage diversity and activation of M-CSF signaling in post-traumatic osteoarthritis"

**Figure S1. Computational subsetting of immune cells from whole synovium scRNA-seq. Related to Figure 1.** (A) Immune cell clusters (2, 5, 8) were computationally subset out of all synovial cells from previously published dataset available at NCBI GEO Accession number GSE211584[1]. (B) Feature plots showing immune or mesenchymal clusters in synovium based on expression of *Ptprc* (encoding CD45) and *Pdgfra*, respectively. (C) Violin plots showing marker genes for each immune cell cluster. (D-F) Total abundance and proportion of major immune cell types, and their subgroupings, in synovium from Sham, 7d ACLR or 28d ACLR mice.

**Figure S2. Recapitulation of published scRNA-seq datasets: Culemann. Related to Figure 2.** (A) UMAP plot of all cell clusters from GSE134420[2]. Cells were sorted CD45+ CD11b+ Ly6G- hindpaw synovial cells from mice subjected to the K/BxN serum transfer model of RA. Clusters are designated as immune or stromal based on the expression of *Ptprc* (encoding CD45) and *Itgam* (encoding CD11b) in (B). The stromal cell cluster (6) was removed, and remaining cells (clusters 0-5) were re-clustered to recapitulate the data found in the originating manuscript (C). Marker genes used to describe each cluster in the original dataset are shown as feature plots (D).

**Figure S3. Recapitulation of published scRNA-seq datasets: Sebastian. Related to Figure 2.** (A) UMAP plot of all cell clusters from GSE200843[3]. Cells were sorted CD45+ cells obtained from a whole knee joint digestion in the ACLR model of mouse PTOA. Clusters are designated as immune, erythrocytes or stromal/vascular based on the expression of *Ptprc* (encoding CD45), *Pdgfra*, *Pecam1* (encoding CD31) and *Hba-a1* (encoding α-globin) in (B). The stromal/vascular and erythrocyte cell clusters (4 and 6) were removed and remaining cells (clusters 0 to 3, 5) were re-clustered to recapitulate the data found in the originating manuscript (C). Marker genes used to describe major cell types in the original dataset are shown as feature plots (D).

**Figure S4. Recapitulation of published scRNA-seq datasets: Muench. Related to Figure 2.** (A) UMAP plot of all cell clusters from GSE184609[4]. Cells were sorted live cells obtained from hindpaw synovia of mice with GPI-induced RA. Clusters are designated as immune, stromal, endothelial, glial, mural, muscle, or proliferating, based on the expression of *Ptprc* (encoding CD45), *Pdgfra*, *Pecam1* (encoding CD31), *Mcam* (encoding CD146), *Acta1* and *Top2a* in (B). All non-immune cell clusters (1, 3, 10 to 14, 18 to 20) were removed and remaining cells (clusters 0, 2, 4 to 9, 15 to 17) were re-clustered to recapitulate the data found in the originating manuscript (C). Marker genes used to describe major cell types in the original dataset are shown as feature plots (D).

**Figure S5. Integration of immune scRNA-seq datasets. Related to Figure 2.** (A) Heatmap of top 5 gene markers for each cell cluster in the integrated immune object of all scRNA-seq datasets. (B) Total abundance of each major immune cell grouping for each dataset. (C) Heatmap of outgoing cell patterns for major immune cell groups generated in CellChat[5]. (D) Outgoing cell signaling pathways that contribute to each pattern in (C).

**Figure S6. Assessment of neutrophils in the whole joint. Related to Figure 2.** (A) Healthy mouse knee joints were subjected to two-step digestion described by Leale *et al*[6], resulting in a soft and hard tissue fraction. Joints were taken from naïve mice (unperfused) or mice perfused with 50 mL PBS to clear removing circulating cells from the vasculature. Plots of live CD45+ cells are shown, and neutrophils (CD45+ CD11b+ Ly6G+) are gated on. Abundance is shown as a proportion of all CD45+ cells in each sample. (B) UMAP plot of all immune cell clusters from all datasets, showing the three neutrophil (Neut-1, 2, 3) clusters. Feature plots of hallmark neutrophil genes are shown below. (C) Top unique genes (with Log2FC and padj) for the three neutrophil clusters were generated by

*FindAllMarkers* then submitted to CIPR[7] analysis using the ImmGen V1 and V2 datasets, to compare the expression signatures of our three neutrophil clusters to pre-sorted published neutrophil gene signatures. Identity score for each pre-sorted cell type is shown. Also see Supplementary Table 2.

**Figure S8. Outgoing signaling patterns from macrophages in PTOA and RA. Related to Figure 3.** (A-B) River plots of outgoing communication patterns of MΦ, monocytes (Mono), and osteoclasts (Osteo) in PTOA (A) or RA (B), generated in CellChat. *Same signals from the same cell type in PTOA and RA. #Same signals but from different cell types in PTOA and RA. $Signals unique to PTOA or RA. (C-D) Top: Heatmaps of outgoing cell patterns for MΦ, monocytes, and osteoclasts in PTOA (C) or RA (D). Bottom: Outgoing cell signaling pathways that contribute to each pattern in PTOA (C) or RA (D).

**Figure S9. Synovial macrophage subsets and trajectories. Related to Figure 4.** (A) To generate an object containing only monocytes and MΦ from our dataset (GSE211584), we first subset all myeloid cell clusters from Seurat into Monocle3, then removed DC, mast cells and granulocytes. (B) Feature plots of classical MΦ/monocyte marker genes, and (C) genes traditionally associated with an M1 or M2 MΦ phenotype. (D) A UMAP plot of all MΦ/monocyte clusters and their designations are provided for reference. n.d.: not detected.

**Figure S10. Stromal-immune crosstalk communication probability ratios in Sham and 7d ACLR. Related to Figure 5.** (A-C) Crosstalk communication probability ratios for multi-directional (stromal ←→ immune) or uni-directional (stromal → immune or immune → stromal) signaling in Sham synovium. A higher ratio indicates a higher likelihood of signaling activity in the specified crosstalk direction. CSF signaling is highlighted in red. (D-F) As above, but for 7d ACLR.

**Figure S11. M-CSF signaling hierarchy to stromal cells. Related to Figure 5.** (A-C) CellChat hierarchy plots for the M-CSF signaling pathway in Sham, 7d ACLR and 28d ACLR, with stromal cells as the receiving cell type in each case. Circles with fill represent cells sending signals, circles without fill represent cells receiving signals, in each condition. No M-CSF signaling was detected in the direction of stromal cells as receivers.

**Figure S12. Expression of key transcription factors across pseudotime from monocytes to macrophages. Related to Figure 6.** (A) RcisTarget analysis was performed to screen promoters of genes enriched in modules 1, 4 and 6 of the monocyte to infiltrating MΦ trajectory. Pseudotime regression plots for the top four transcription factors predicted by RcisTarget to regulate enriched module genes are shown.

### Fig. S1. Computational subsetting of immune cells from whole synovium scRNA-seq.



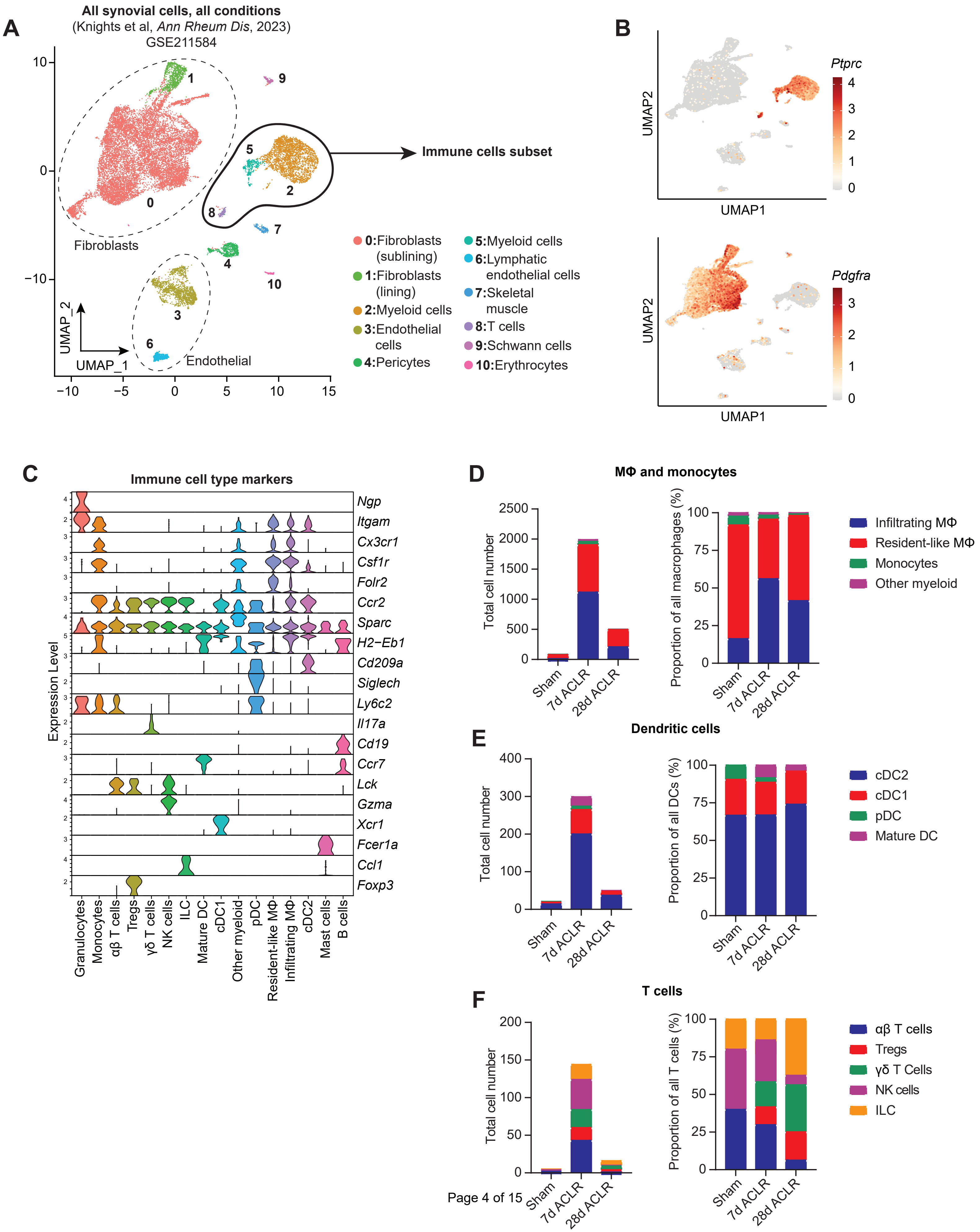

### Fig. S2. Recapitulation of published scRNA-seq datasets: Culemann.



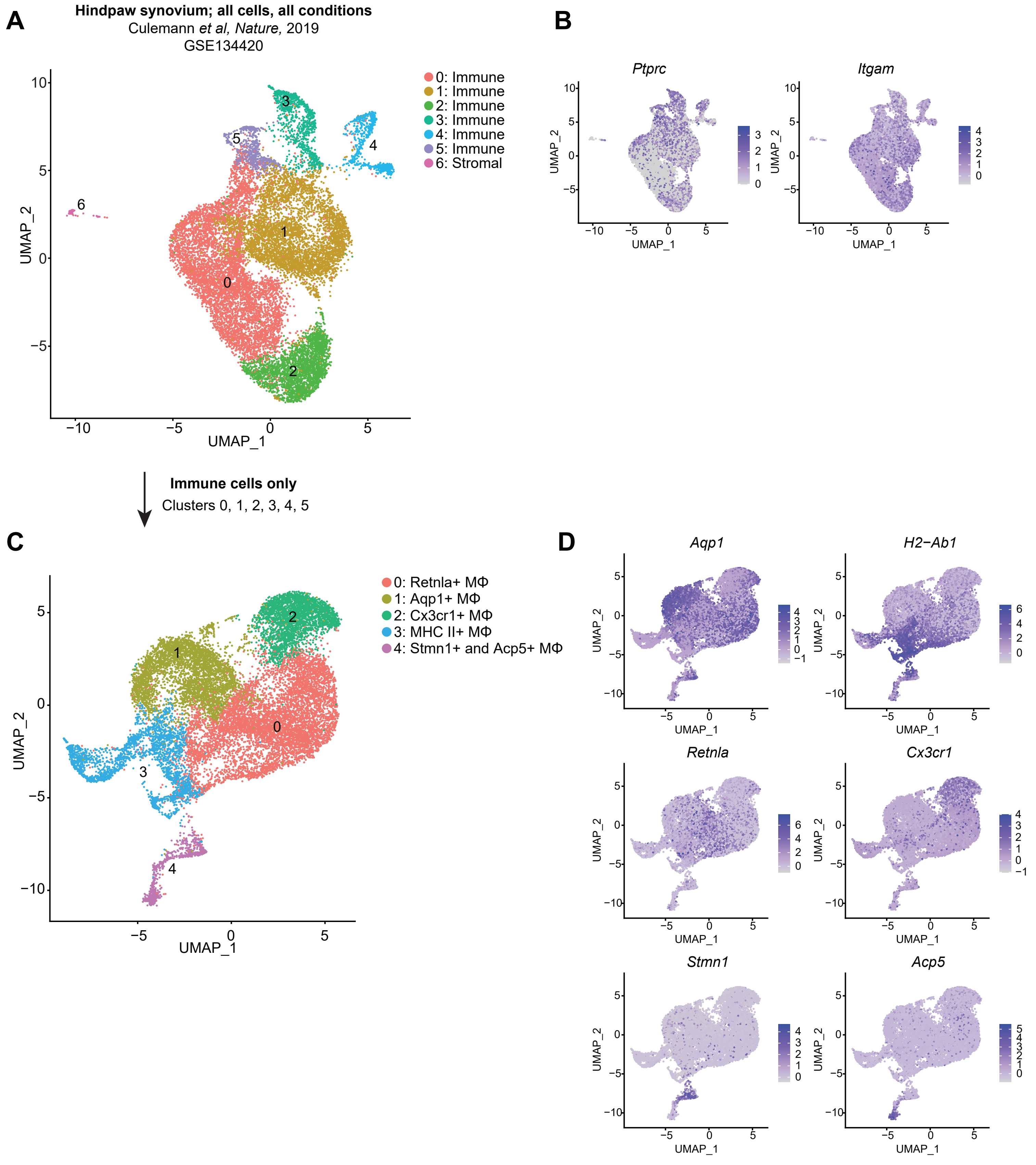

### Fig. S3. Recapitulation of published scRNA-seq datasets: Sebastian.

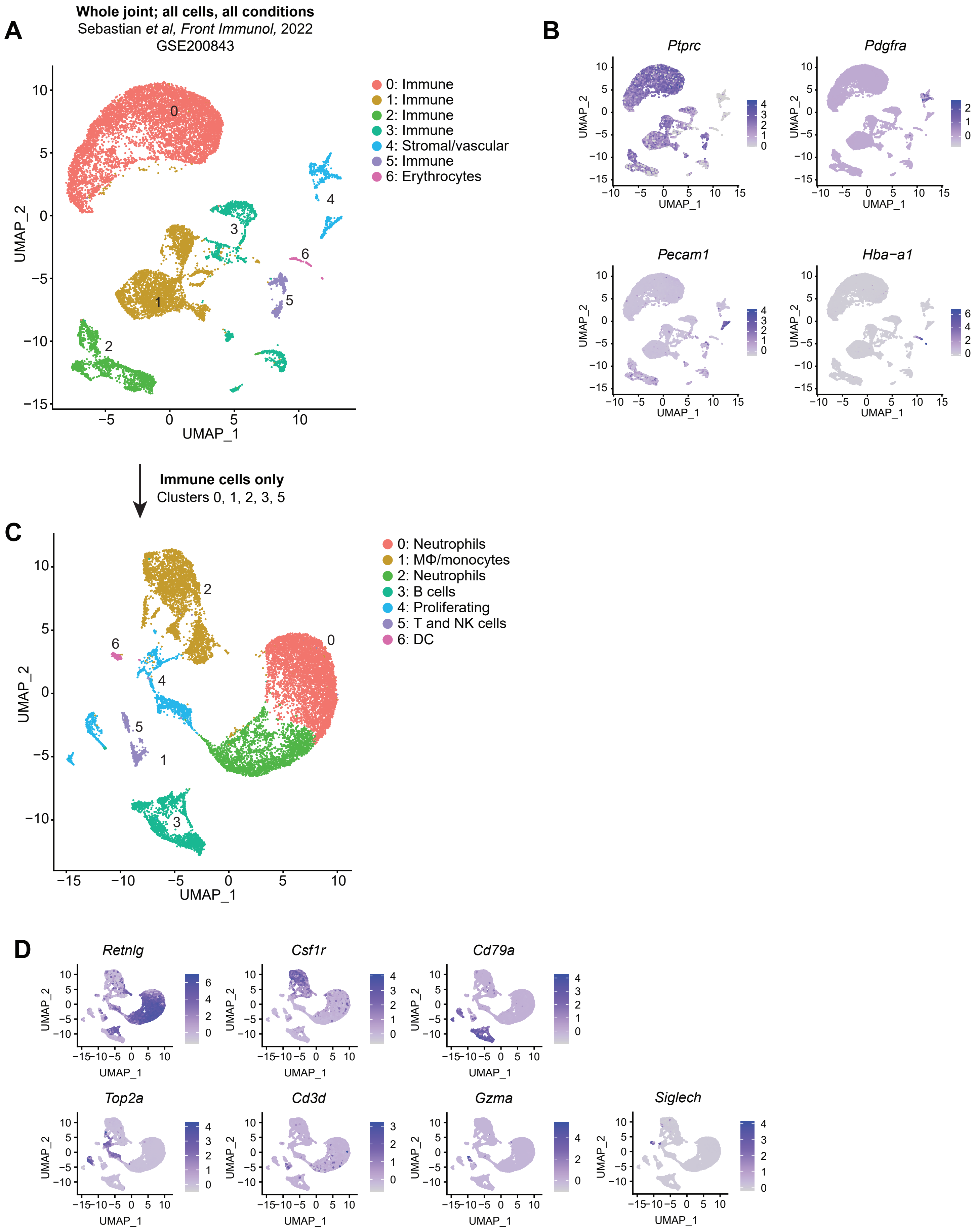

### Fig. S4. Recapitulation of published scRNA-seq datasets: Muench.



**Hindpaw synovium; all cells, all conditions**
Muench *et al, J Immunol,* 2022
GSE184609

**A**

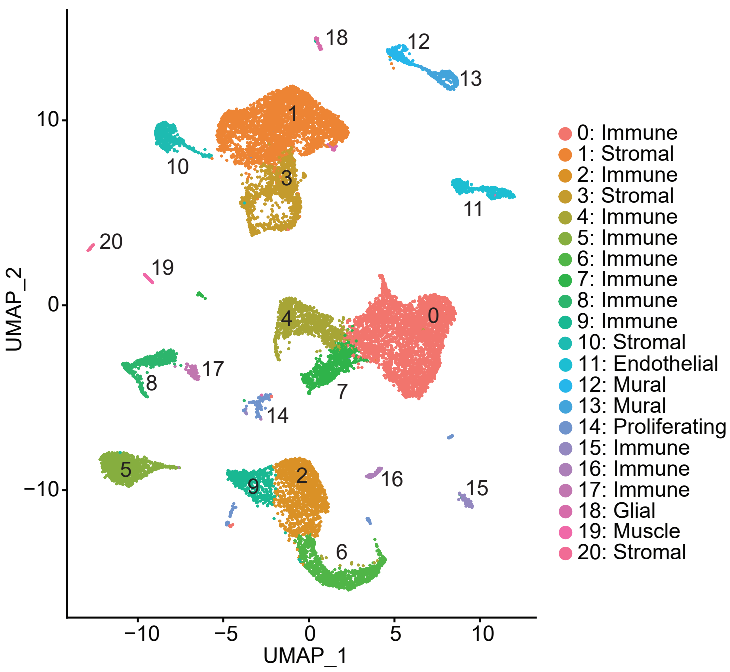


**B**

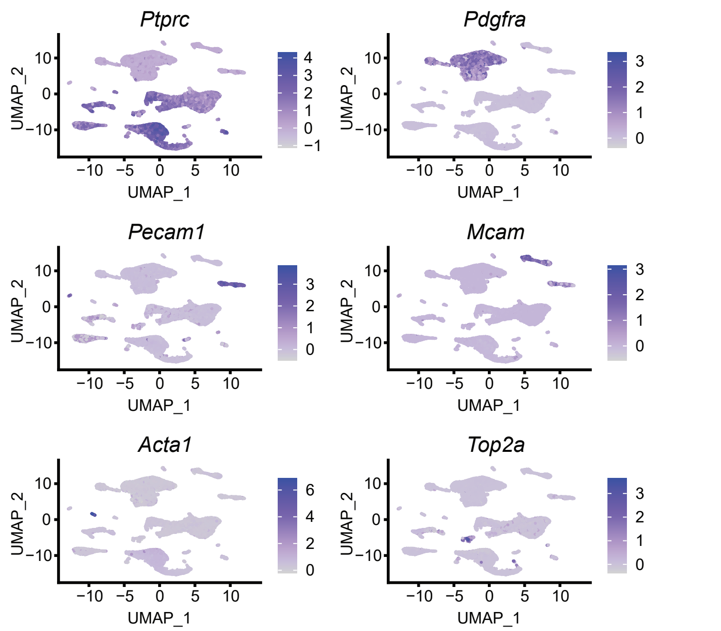


**C**

**Immune cells only**
Clusters 0, 2, 4, 5, 6, 7, 8, 9, 11, 15, 16, 17

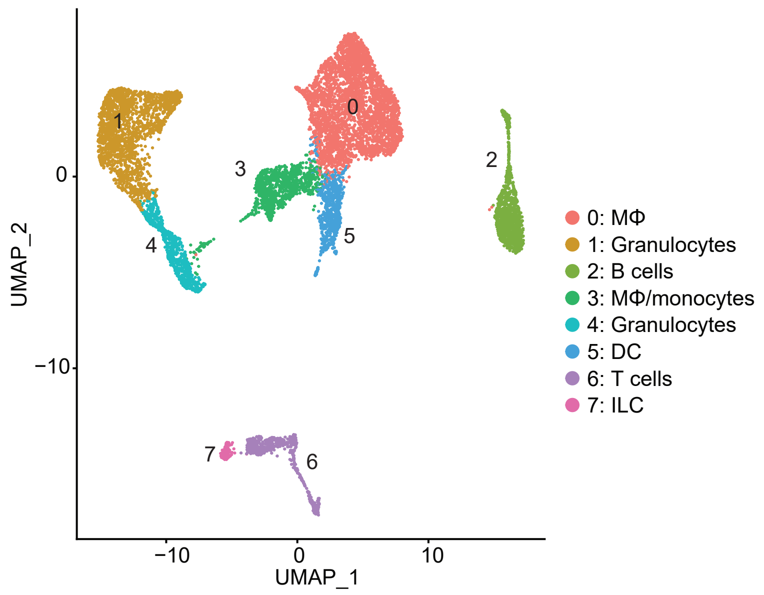


**D**

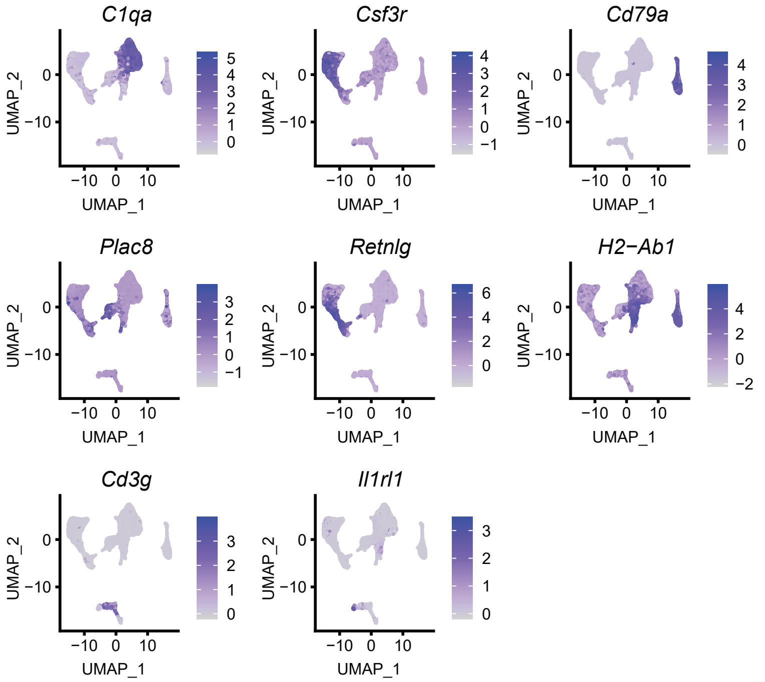

### Fig. S5. Integration of immune scRNA-seq datasets.

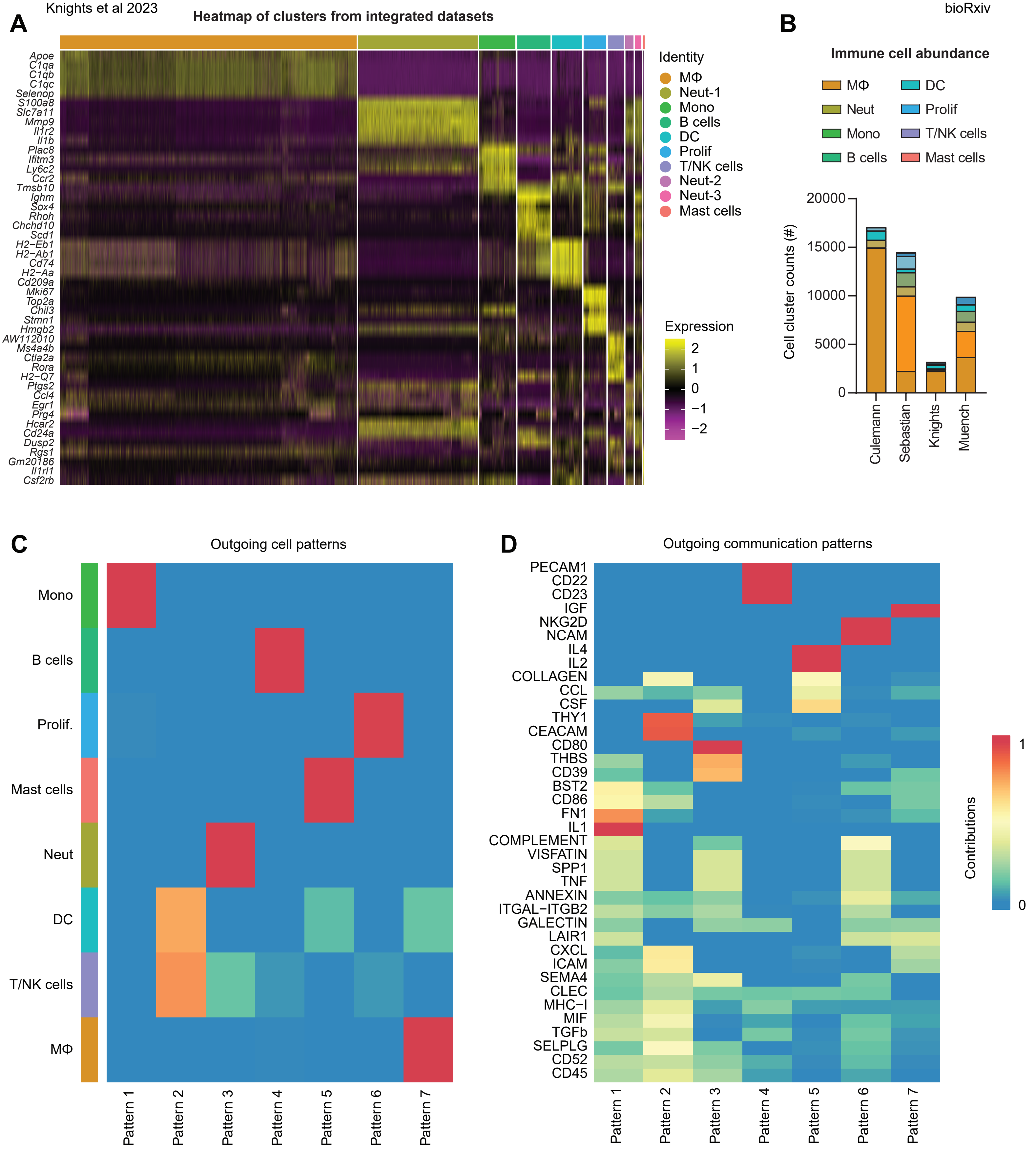

### Fig. S6. Assessment of neutrophils in the whole joint.



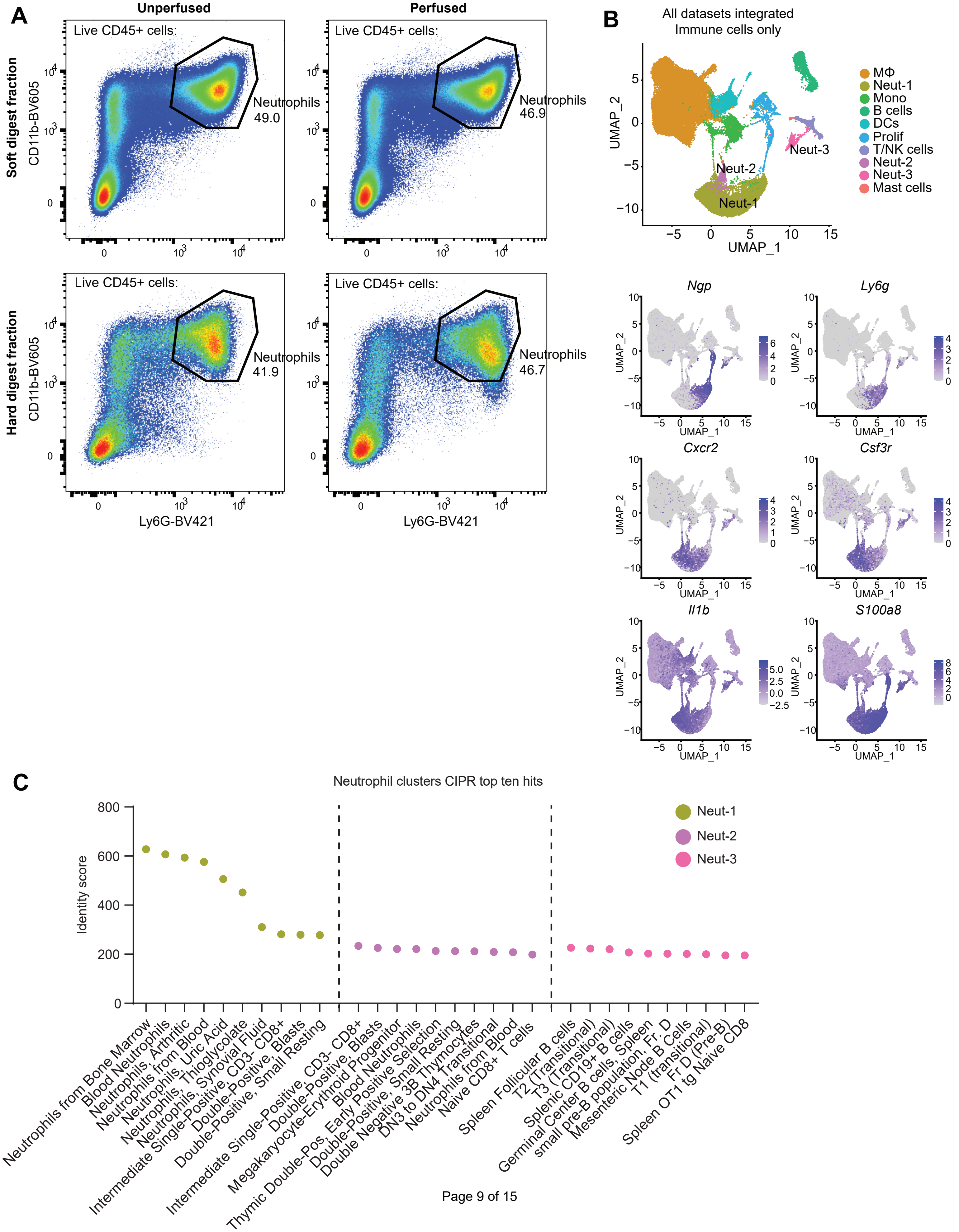

#### Fig. S7. Macrophages in OA and RA





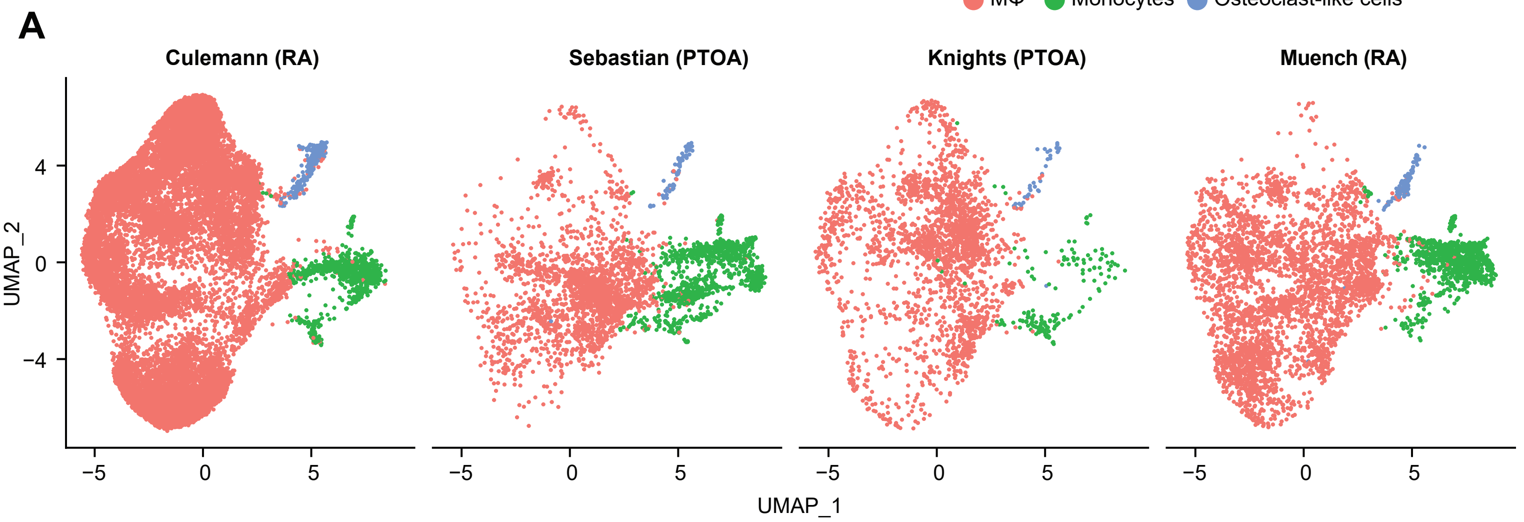

### Fig. S8. Outgoing signaling patterns from macrophages in PTOA and RA

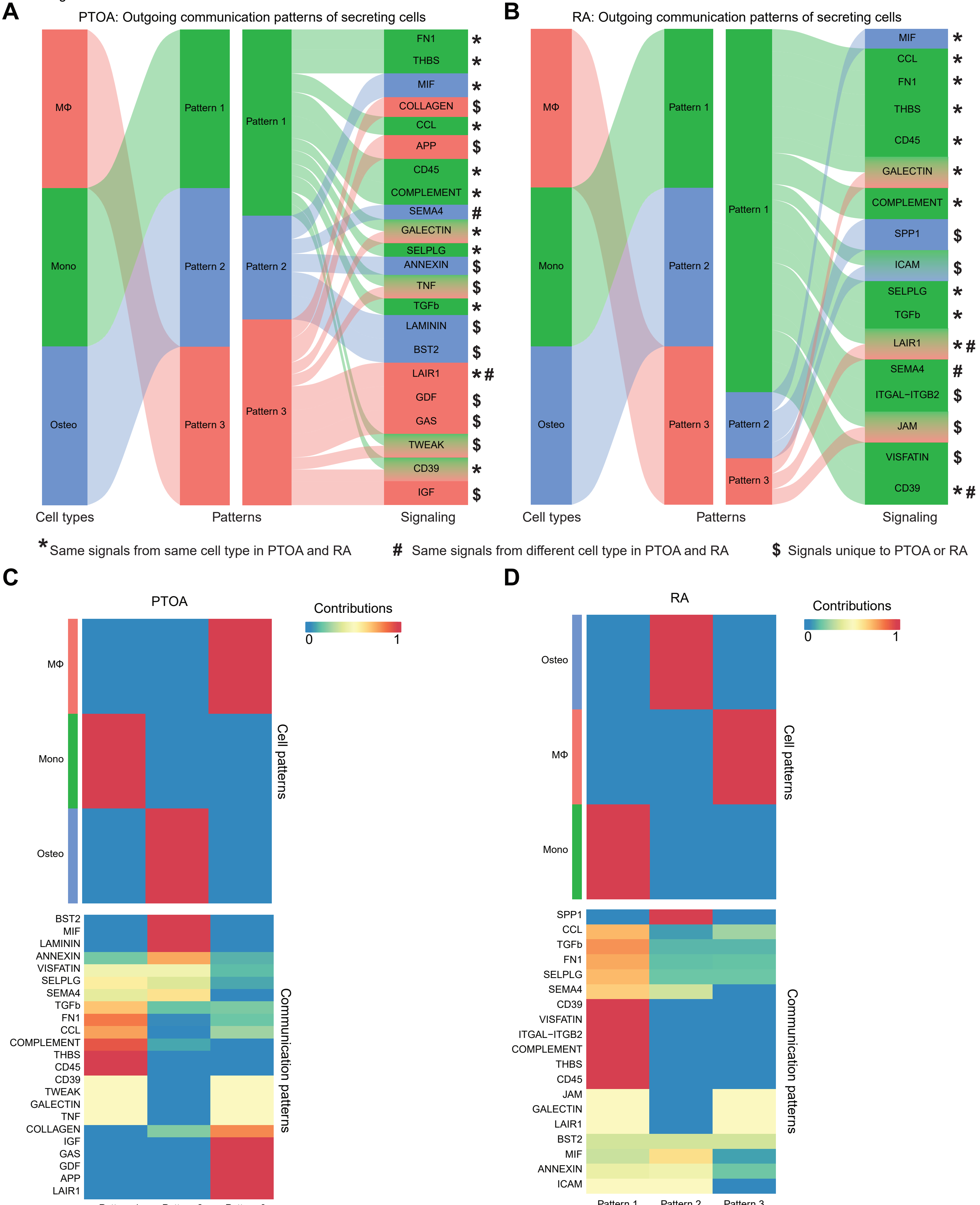

**Fig. S9. Synovial macrophage subsets and trajectories.**

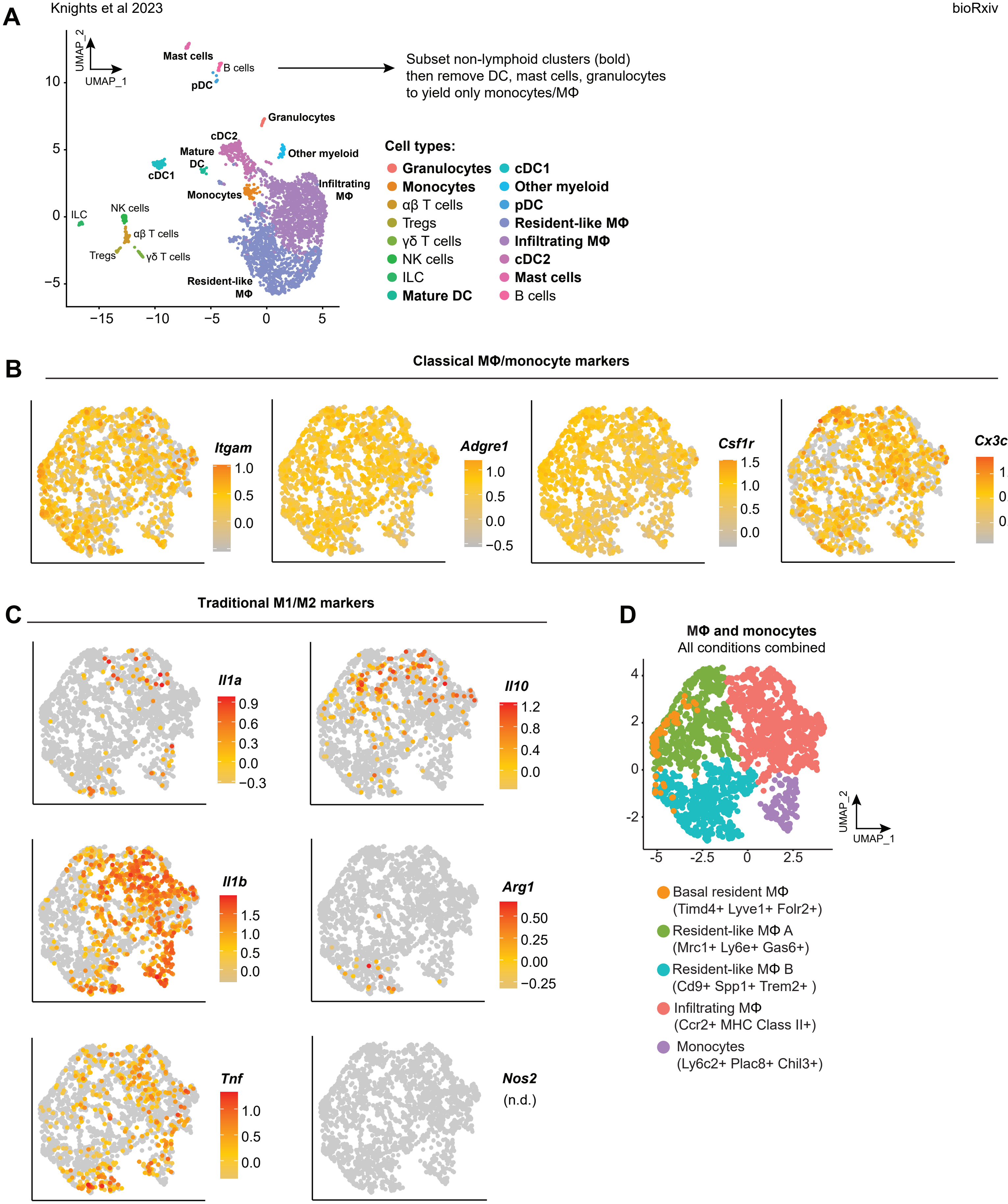

### Fig. S10. Stromal-immune crosstalk communication probability ratios in Sham and 7d ACLR

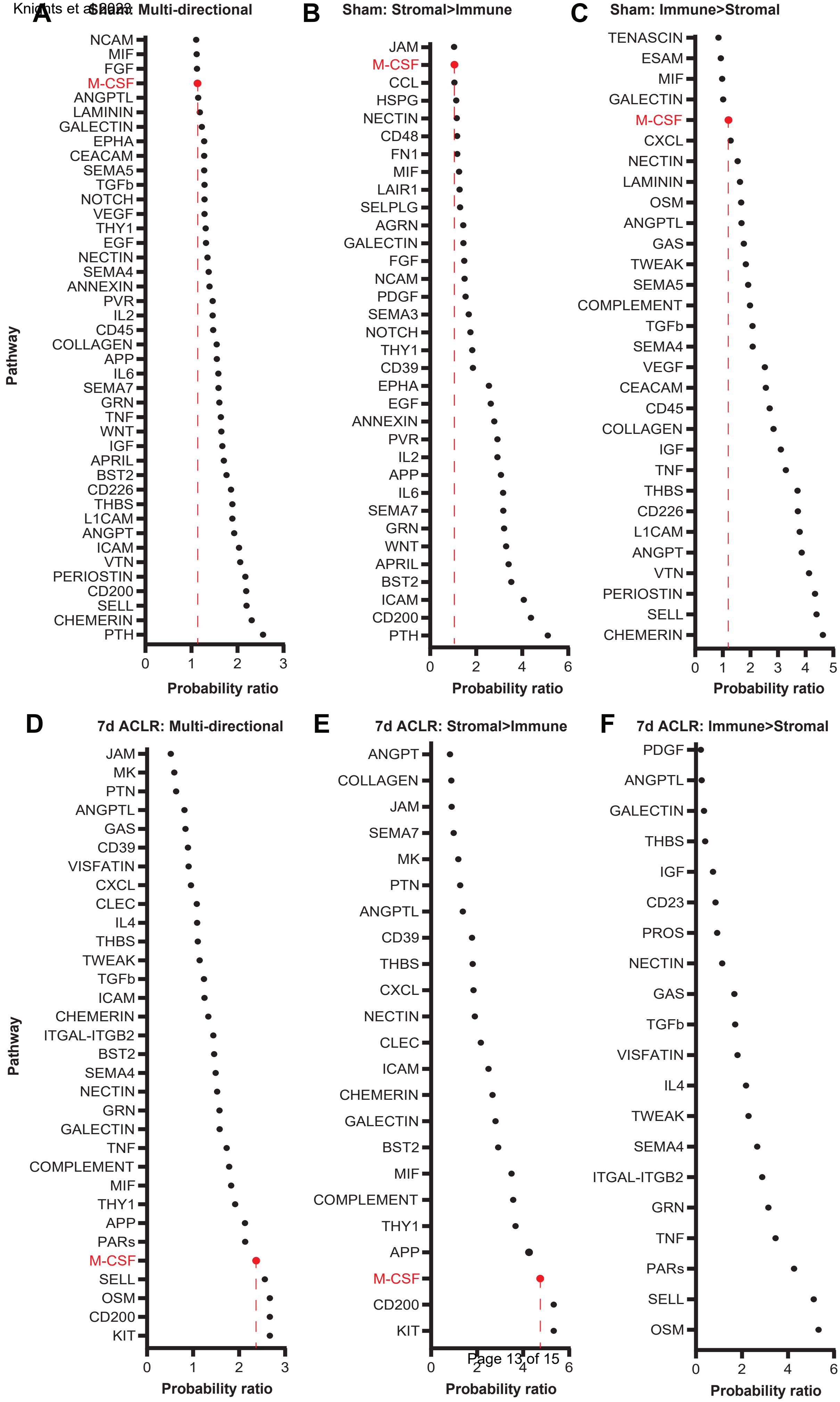

#### Fig. S11. M-CSF signaling hierarchy to stromal cells



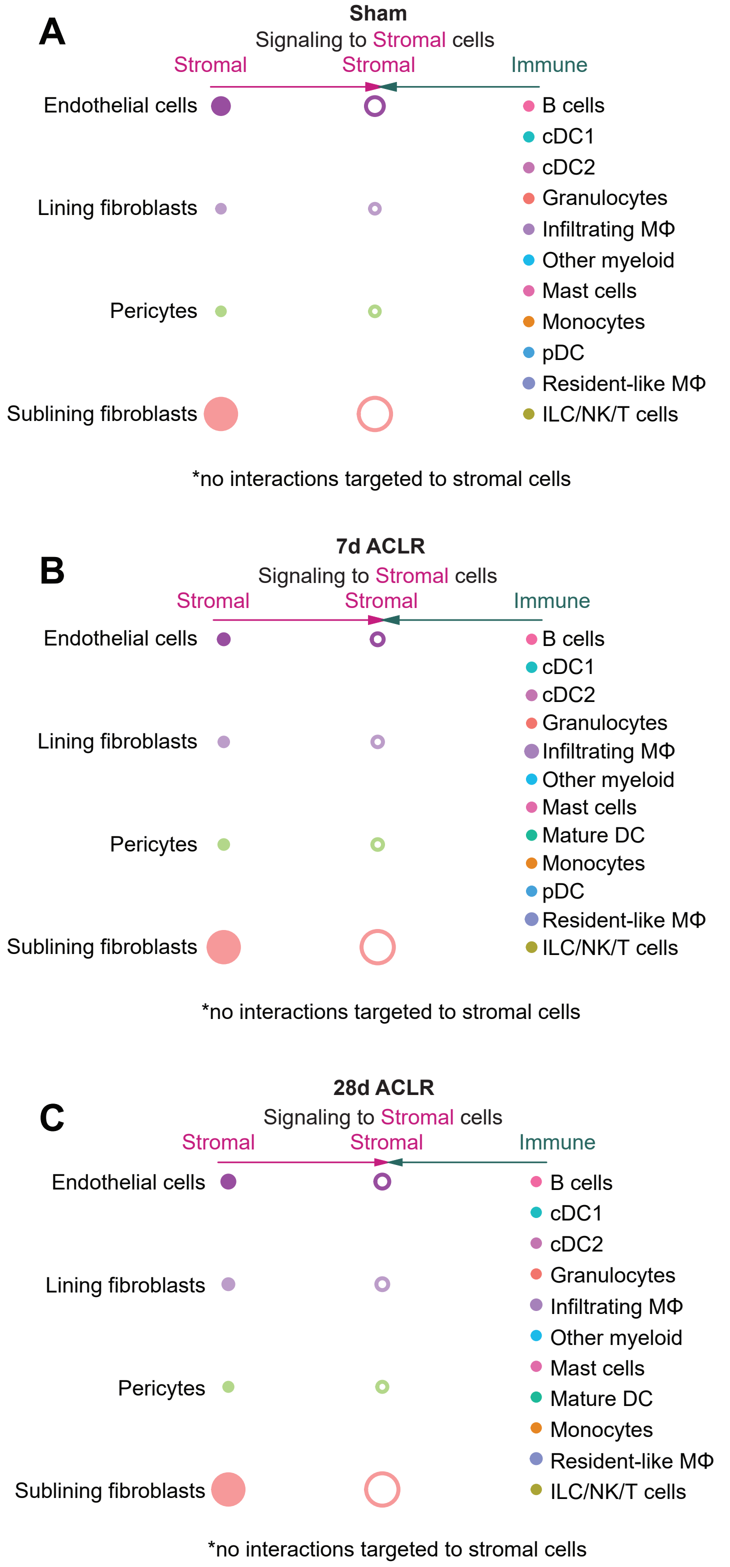

### Fig. S12. Expression of key transcription factors across pseudotime from monocytes to macrophages.



A

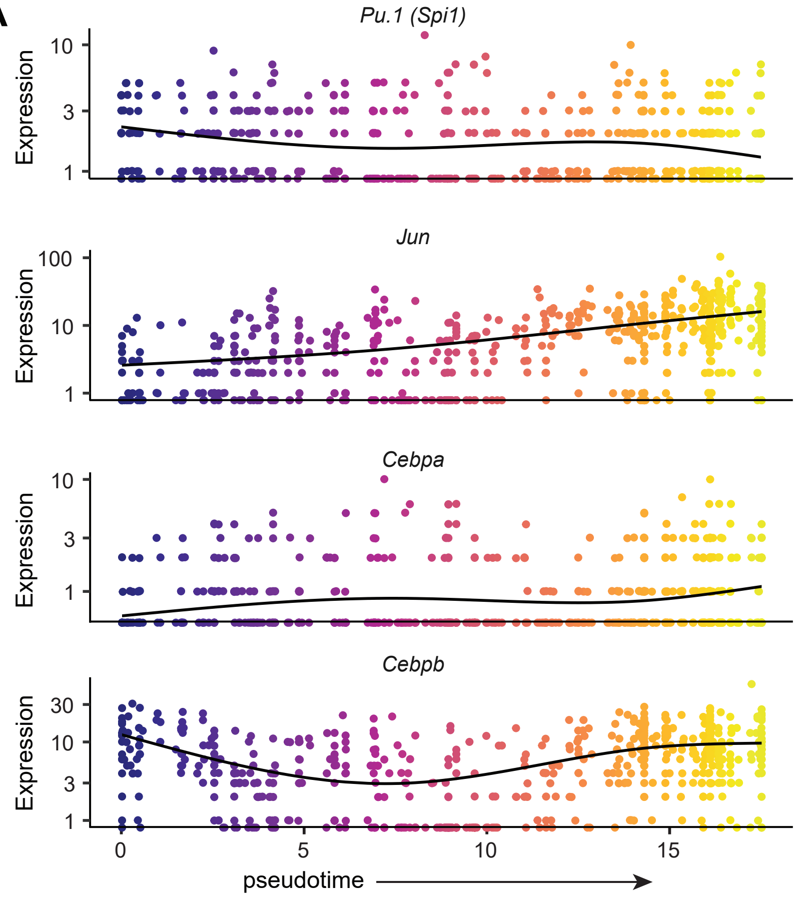
