## Supplementary table 1 for "Synovial macrophage diversity and activation of M-CSF signaling in post-traumatic osteoarthritis"

**Supplementary Table 1. Flow cytometry antibodies.**

| **Antibody name** | **Clone** | **Vendor** | **RRID** | **Dilution** |
| --- | --- | --- | --- | --- |
| Anti-mouse CD3-APC/Fire750 | 17A2 | Biolegend #100247 | AB_2572117 | 1:100 |
| Anti-mouse CD11b-BV605 | M170 | Biolegend #101237 | AB_11126744 | 1:400 |
| Anti-mouse CD11c-PE/Dazzle594 | N418 | Biolegend #117347 | AB_2563654 | 1:200 |
| Anti-mouse CD19-FITC | 1D3/CD19 | Biolegend #152404 | AB_2629813 | 1:100 |
| Anti-mouse CD45-BV650 | 30-F11 | Biolegend #103151 | AB_2565884 | 1:400 |
| Anti-mouse F4/80-APC/R700 | T45-2342 | BD Horizon #565787 | AB_2869711 | 1:400 |
| Anti-mouse FceRIa-PE/Cy7 | MAR-1 | Biolegend #134318 | AB_10640122 | 1:100 |
| Anti-mouse Ly6G-BV421 | 1A8 | Biolegend #127627 | AB_10897944 | 1:200 |
| Anti-mouse MHC Class II-BV421 | M5/114.15.2 | Biolegend #107632 | AB_2650896 | 1:200 |
| Anti-mouse NK1.1-BV510 | PK136 | Biolegend #108737 | AB_2562216 | 1:200 |
