## Supplementary table 2 for "Synovial macrophage diversity and activation of M-CSF signaling in post-traumatic osteoarthritis"

**Supplementary Table 2. Top ten CIPR hits for each neutrophil cluster from the integrated object of all immune cells, all datasets.**

| **cluster** | **reference_id** | **identity_score** | **long_name** | **reference_cell_type** | **description** |
| --- | --- | --- | --- | --- | --- |
| Neut-1 | GN.BM | 627.7323155 | Neutrophils from Bone Marrow | Granulocyte | Phenotype markers: CD11b+ Ly6-G+ // Exclusion markers: NA // Tissue: femur // Age: 6 weeks // Gender: male // Genetic background: C57BL/6J // ImmGen release: v1 |
| Neut-1 | GN.Bl.v2 | 606.899114 | Blood Neutrophils | Granulocyte | Phenotype markers: SSChi Ly6Ghi // Exclusion markers: B220, CD4 // Tissue: Blood // Age: 7 Weeks // Gender: NA // Genetic background: C57Bl/6J // ImmGen release: v2 |
| Neut-1 | GN.Arth.BM | 593.463945 | Neutrophils, Arthritic | Granulocyte | Phenotype markers: CD11b+ Ly6-G+ // Exclusion markers: NA // Tissue: Bone marrow // Age: 6 weeks // Gender: male // Genetic background: C57BL/6J // ImmGen release: v1 |
| Neut-1 | GN.Bl | 576.3919565 | Neutrophils from Blood | Granulocyte | Phenotype markers: CD11b+ Ly6-G+ // Exclusion markers: NA // Tissue: blood // Age: 6 weeks // Gender: male // Genetic background: C57BL/6J // ImmGen release: v1 |
| Neut-1 | GN.UrAc.PC | 506.609048 | Neutrophils, Uric Acid | Granulocyte | Phenotype markers: CD11b+ Ly6-G+ // Exclusion markers: NA // Tissue: Peritoneal Cavity // Age: 6 weeks // Gender: male // Genetic background: C57BL/6J // ImmGen release: v1 |
| Neut-1 | GN.Thio.PC | 451.6901174 | Neutrophils, Thioglycolate | Granulocyte | Phenotype markers: CD11b+ Ly6-G+ // Exclusion markers: NA // Tissue: Peritoneal Cavity // Age: 6 weeks // Gender: male // Genetic background: C57BL/6J // ImmGen release: v1 |
| Neut-1 | GN.Arth.SynF | 310.0830606 | Neutrophils, Synovial Fluid | Granulocyte | Phenotype markers: CD11b+ Ly6-G+ // Exclusion markers: NA // Tissue: Synovial Fluid // Age: 6 weeks // Gender: male // Genetic background: C57BL/6J // ImmGen release: v1 |
| Neut-1 | T.ISP.Th | 280.87816 | Intermediate Single-Positive, CD3- CD8+ | Pre-T cell | Phenotype markers: 4- 8+ TCR-/lo 24hi // Exclusion markers: 11b- 11c- 19- 49b- Gr1- TER119- // Tissue: thymus // Age: 6 weeks // Gender: male // Genetic background: C57BL/6J // ImmGen release: v1 |
| Neut-1 | T.DPbl.Th | 278.7712978 | Double-Positive, Blasts | Pre-T cell | Phenotype markers: 4+ 8+ TCR-/lo FSChi // Exclusion markers: 11b- 11c- 19- 49b- Gr1- TER119- // Tissue: thymus // Age: 6 weeks // Gender: male // Genetic background: C57BL/6J // ImmGen release: v1 |
| Neut-1 | T.DPsm.Th | 277.5728519 | Double-Positive, Small Resting | Pre-T cell | Phenotype markers: 4+ 8+ TCR-/lo FSClo // Exclusion markers: 11b- 11c- 19- 49b- Gr1- TER119- // Tissue: thymus // Age: 6 weeks // Gender: male // Genetic background: C57BL/6J // ImmGen release: v1 |
| Neut-2 | T.ISP.Th | 233.8869173 | Intermediate Single-Positive, CD3- CD8+ | Pre-T cell | Phenotype markers: 4- 8+ TCR-/lo 24hi // Exclusion markers: 11b- 11c- 19- 49b- Gr1- TER119- // Tissue: thymus // Age: 6 weeks // Gender: male // Genetic background: C57BL/6J // ImmGen release: v1 |
| Neut-2 | T.DPbl.Th | 225.7463305 | Double-Positive, Blasts | Pre-T cell | Phenotype markers: 4+ 8+ TCR-/lo FSChi // Exclusion markers: 11b- 11c- 19- 49b- Gr1- TER119- // Tissue: thymus // Age: 6 weeks // Gender: male // Genetic background: C57BL/6J // ImmGen release: v1 |
| Neut-2 | SC.MEP.BM | 221.0333307 | Megakaryocyte-Erythroid Progenitor | Stem-Progenitor | Phenotype markers: Lineage- Kit+ Sca1- CD34- FcgR-/low // Exclusion markers: PI // Tissue: Bone marrow // Age: 9 weeks // Gender: male // Genetic background: C57BL/6J // ImmGen release: v1 |
| Neut-2 | GN.Bl.v2 | 220.8742952 | Blood Neutrophils | Granulocyte | Phenotype markers: SSChi Ly6Ghi // Exclusion markers: B220, CD4 // Tissue: Blood // Age: 7 Weeks // Gender: NA // Genetic background: C57Bl/6J // ImmGen release: v2 |
| Neut-2 | T.DP.Th.v2 | 213.0242692 | Thymic Double-Positive, Early Positive Selection | T cell | Phenotype markers: CD4+ CD8+ CD69- // Exclusion markers: NA // Tissue: Thymus // Age: 6 Weeks // Gender: NA // Genetic background: NA // ImmGen release: v2 |
| Neut-2 | T.DPsm.Th | 212.332587 | Double-Positive, Small Resting | Pre-T cell | Phenotype markers: 4+ 8+ TCR-/lo FSClo // Exclusion markers: 11b- 11c- 19- 49b- Gr1- TER119- // Tissue: thymus // Age: 6 weeks // Gender: male // Genetic background: C57BL/6J // ImmGen release: v1 |
| Neut-2 | preT.DN3B.Th | 211.7100238 | Double Negative 3B Thymocytes | Pre-T cell | Phenotype markers: Lin-/lo CD25hi CD44- CD28+ // Exclusion markers: Lin= CD4, CD8a, TCRb, TCRgd, CD11b, CD11c, NK1.1, Gr1, CD19, Ter119 // Tissue: thymus // Age: 6 weeks // Gender: male // Genetic background: C57BL/6J // ImmGen release: v1 |
| Neut-2 | preT.DN3-4.Th | 208.7467353 | DN3 to DN4 Transitional | Pre-T cell | Phenotype markers: Lin-/lo CD25int CD44- CD28+ // Exclusion markers: Lin= CD4, CD8a, TCRb, TCRgd, CD11b, CD11c, NK1.1, Gr1, CD19, Ter119 // Tissue: thymus // Age: 6 weeks // Gender: male // Genetic background: C57BL/6J // ImmGen release: v1 |
| Neut-2 | GN.Bl | 207.3685017 | Neutrophils from Blood | Granulocyte | Phenotype markers: CD11b+ Ly6-G+ // Exclusion markers: NA // Tissue: blood // Age: 6 weeks // Gender: male // Genetic background: C57BL/6J // ImmGen release: v1 |
| Neut-2 | CD8.5h.LN | 198.0018699 | Naive CD8+ T cells 5 hours after in vitro aCD3-aCD28 stimulation | T cell | Phenotype markers: CD8+ CD4- TCR+ CD25- CD62Lhi CD44 Lo // Exclusion markers: CD11c/19/25/49b/Gr1/Ter119- // Tissue: Lymph Node // Age: 6 Weeks OLD // Gender: NA // Genetic background: C57BL/6 // ImmGen release: v2 |
| Neut-3 | B.Fo.Sp | 226.2901762 | Spleen Follicular B cells | B cell | Phenotype markers: IgD+ IgM+ CD45R+ CD24+ CD19+ AA4.1- CD23+ // Exclusion markers: CD3 Ly6c Ter119 Gr-1 Mac1 // Tissue: spleen // Age: 7 weeks // Gender: male // Genetic background: C57BL/6J // ImmGen release: v1 |
| Neut-3 | B.T2.Sp | 223.1088855 | T2 (Transitional) | B cell | Phenotype markers: IgD+ IgM+ CD45R+ CD24+ CD19+ AA4.1+ CD23+ // Exclusion markers: CD3 Ly6c Ter119 Gr-1 Mac1 // Tissue: spleen // Age: 7 weeks // Gender: male // Genetic background: C57BL/6J // ImmGen release: v1 |
| Neut-3 | B.T3.Sp | 220.6418424 | T3 (Transitional) | B cell | Phenotype markers: IgD+ IgM+ CD45R+ CD24+ CD19+ AA4.1+ CD23+ // Exclusion markers: CD3 Ly6c Ter119 Gr-1 Mac1 // Tissue: spleen // Age: 7 weeks // Gender: male // Genetic background: C57BL/6J // ImmGen release: v1 |
| Neut-3 | CD19Control | 207.2535992 | Splenic CD19+ B cells | B cell | Phenotype markers: 19+ 4- 8- // Exclusion markers: NA // Tissue: spleen // Age: 6 weeks // Gender: male // Genetic background: C57BL/6J // ImmGen release: v1 |
| Neut-3 | B.GC.Sp | 202.1446785 | Germinal Center B cells, Spleen | B cell | Phenotype markers: CD19+ IgM+ IgD- GL7+ PNA+ // Exclusion markers: Ter119 Ly6c CD3 CD11b Gr-1 // Tissue: spleen // Age: 10 weeks // Gender: male // Genetic background: C57BL/6J // ImmGen release: v1 |
| Neut-3 | preB.FrD.BM | 201.8734309 | small pre-B population, Fr. D | Pre-B cell | Phenotype markers: CD19+ IgM- CD45R+ CD43- // Exclusion markers: CD3 Ter119 Ly6c Cd11b Gr-1 // Tissue: Bone marrow // Age: 6 weeks // Gender: male // Genetic background: C57BL/6J // ImmGen release: v1 |
| Neut-3 | B.Fo.MLN | 201.3055125 | Mesenteric Node B Cells | B cell | Phenotype markers: CD19+ CD45R+ CD23+ CD21/35+ // Exclusion markers: CD3 TER119 GR1 Ly6c AA4.1 CD24 IgM CD43 // Tissue: Mesenteric LN // Age: 8 weeks // Gender: male // Genetic background: C57BL/6J // ImmGen release: v1 |
| Neut-3 | B.T1.Sp | 199.3544315 | T1 (transitional) | B cell | Phenotype markers: IgD+ IgM+ CD45R+ CD24+ CD19+ AA4.1+ CD23- // Exclusion markers: CD3 Ly6c Ter119 Gr-1 Mac1 // Tissue: spleen // Age: 7 weeks // Gender: male // Genetic background: C57BL/6J // ImmGen release: v1 |
| Neut-3 | preB.FrD.FL | 195.1916109 | Fr D (Pre-B) | Pre-B cell | Phenotype markers: AA4.1+ IgM- CD19+ CD43- CD24+ // Exclusion markers: Ter119 Ly6c CD11b Gr-1 CD3 // Tissue: Fetal Liver // Age: 18 days // Gender: both (pooled) // Genetic background: C57BL/6J // ImmGen release: v1 |
| Neut-3 | T.8Nve.Sp.OT1 | 194.7049542 | Spleen OT1 tg Naive CD8 | T cell | Phenotype markers: CD8+ CD45.1+ // Exclusion markers: B220- CD45.2- Gr-1- NK1.1- CD4- MHCII- P.I.- // Tissue: spleen // Age: 6 weeks // Gender: male // Genetic background: C57BL/6J // ImmGen release: v1 |
